# Profiling histone post-translational modifications to identify signatures of epigenetic drug response in T-cell acute lymphoblastic leukemia

**DOI:** 10.1101/2025.09.01.673463

**Authors:** Laura Corveleyn, Lien Provez, Osman Satilmis, Nina Refhagen, Mattias Landfors, Wouter Sleeckx, Beatrice Lintermans, Amélie De Maesschalck, Rishi S. Kotecha, Barbara De Moerloose, Tim Lammens, Dieter Deforce, Sofie Degerman, Steven Goossens, Pieter Van Vlierberghe, Maarten Dhaenens

## Abstract

Epigenetic modifications are dynamic and reversible, making them attractive targets for therapeutic intervention in cancer. Although several epigenetic drugs (epidrugs) have been clinically approved, their application in T-cell acute lymphoblastic leukemia (T-ALL) remains limited, and predictive biomarkers of response are lacking. Here, we present a mass spectrometry (MS)-based pharmacoepigenetic approach to profile histone post-translational modifications (hPTMs) to identify signatures associated with epidrug sensitivity in T-ALL. Baseline hPTM landscapes were previously established by our group for 21 T-ALL cell lines using liquid chromatography–tandem mass spectrometry (LC–MS/MS). Here, we treated these cell lines with a panel of nine epidrugs including anthracyclines, histone deacetylase inhibitors, and DNA methyltransferase inhibitors. Correlation of cell viability data with hPTM levels revealed distinct hPTM signatures linked to sensitivity for each drug class. These signatures were subsequently evaluated in T-ALL patient-derived xenograft (PDX) models. However, we our analysis revealed substantial discepancies in hPTM sensitivity signatures compared to those observed in vitro. Co-variation network analysis highlighted divergence in hPTM-hPTM correlation between the two models, underscoring limitations of cell lines for modeling dynamic epigenetic regulation in vivo. Our findings establish a framework for MS-based hPTM profiling in T-ALL and emphasize the importance of model selection in developing predictive epigenetic biomarkers.

**Graphical abstract:** 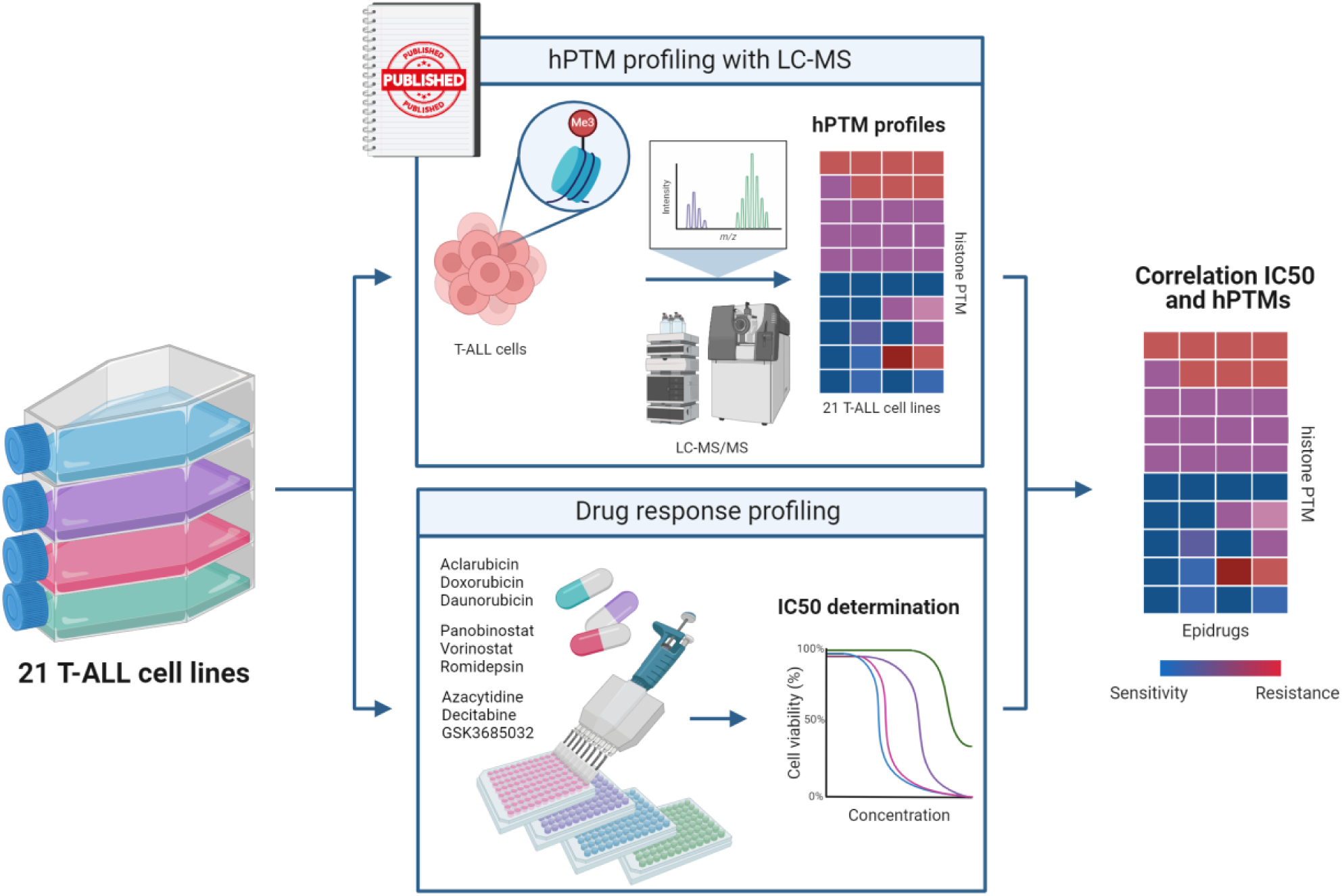

Global hPTM profiling of 21 T-ALL cell lines was performed using LC-MS/MS, as previously published by Provez et al. In parallel, the 21 cell lines were treated with a dilution series of nine epidrugs, categorized into three distinct classes, to determine their IC50 values. Finally, Spearman correlation analysis was performed to assess the relationship between hPTM levels and drug sensitivity. Figure created with Biorender.com.

## Introduction

Resistance to cancer therapy remains one of the key challenges of modern oncology. The identification of predictive biomarkers could provide information on the risk-to-benefit ratio of a given drug and thereby allow personalized treatment. In the last few decades, major advances have been made in the field of pharmacogenomics, which aims at understanding how genetic variations affect drug efficacy and toxicity^1^. However, genetic variations account for only a subset of patient-specific factors that influence variability in drug response. The remainder is explained by a variety of environmental and lifestyle factors, such as age, sex, ethnicity, diet, physical activity, and concomitant drug use. These factors can in turn affect gene expression patterns by modulating gene regulation mechanisms, which is called epigenetics. The rapidly growing interest in the integration of epigenetics into the field of pharmaceutical sciences has led to the recently coined term “pharmacoepigenetics”, i.e. the study of the non-heritable bases for interindividual variability in drug response^2^.

The most extensively studied and clinically exploited epigenetic modification is DNA methylation, which is generally associated with gene silencing, i.e. interrupting the transcription of certain genes^3^. A second, less well studied epigenetic template are the evolutionary conserved histone proteins that are associated with DNA in the nucleus to form the structural units of chromatin. Several dynamic post-translational modifications (PTMs) on these proteins together make up the “histone code”. Histone PTMs (hPTMs) modulate the accessibility of DNA to transcription factors, leading to alterations in gene expression. These alterations can be attributed to either the chemical properties of hPTMs or to their ability to recruit modifier enzymes and binding proteins^4^.

Unlike genetic mutations, which are generally irreversible, epigenetic modifications are dynamic and can be reversed, which makes them ideal drug targets. Notably, drugs that regulate epigenetic signatures associated with particular disease patterns, termed epidrugs, have already found their way into the clinic, including DNA methyltransferase (DNMT) inhibitors and histone deacetylase (HDAC) inhibitors^2^. Interestingly, most of these new epidrugs are approved in the EU for the indications of rare blood cancers^5^. However, they are not yet part of standard treatment protocols and are mainly used under compassionate or medical-need programs for relapsed or refractory cases after standard therapies have failed.

T-cell acute lymphoblastic leukemia (T-ALL) is an aggressive hematologic malignancy proceeding from the aberrant transformation of immature T-cell progenitors^6^. T-ALL occurs more frequently in males then females and is predominantly identified in children and adolescents. The standard of care for T-ALL consists of high-dose multi-agent chemotherapy potentially followed by hematopoietic stem cell transplantation (HSCT). While modern treatment regimens cure more than 85 % of children and approximately half of adults, a significant proportion of patients still face poor outcomes^7–10^. In addition, outcomes for patients with primary refractory disease or relapse remain extremely low. Moreover, such intensified multimodal chemotherapy protocols are associated with substantial short- and long-term side-effects, and outcomes for patients with primary refractory disease or relapse remain extremely low. Therefore, there is a compelling need for predictive (epi)genetic biomarkers and novel therapies.

To date, most pharmacoepigenetic research has relied on DNA methylation or antibody-based assays (e.g. ChipSeq, Western Blot) for hPTM analysis, which are limited to the detection of one PTM per experiment^11–13^. However, due to the combinatorial nature of the histone language, targeting single PTMs provides little power for predicting drug sensitivity. Therefore, a bird’s eye view of the histone code is required to detect the hundreds of different hPTMs simultaneously and map their complex interplay in great detail. Mass spectrometry (MS) is currently the only technique capable of examining the complex dynamics of hPTMs in an untargeted way^14^. A comprehensive assay studying the histone code as a tool for facilitating personalized medicine is currently lacking.

We propose to fill this void by developing the first MS-based pharmacoepigenetic assay, targeting hPTMs, for predicting epidrug response in T-ALL. Previously, we established a comprehensive histone T-ALL atlas by measuring the baseline histone PTM profile of 21 different T-ALL cell lines using LS-MS/MS^15^. Here, we treated these 21 T-ALL cell lines with a dilution series of nine epidrugs, subdivided in three classes: anthracyclines, histone deacetylase (HDAC) inhibitors and DNA methyltransferase (DNMT) inhibitors. By linking dose-response curves to baseline hPTM profiles, hPTM signatures arose that are indicative of sensitivity to each compound. Subsequently, the validity of these signatures was tested ex vivo in patient-derived xenografts (PDXs) cell cultures. However, this revealed significant discrepancies between in vitro cell lines and ex vivo patient derived cell culture conditions. Co-variation analysis, illustrated through correlograms, demonstrates that the co-regulation of long-distance histone post-translational modifications (hPTMs) varies drastically between these models. This highlights that cell lines may not accurately represent the complex epigenetic interactions occurring in vivo, further questioning their reliability as a model for studying dynamic epigenetic regulation.

## Results

### hPTM signatures can be correlated to epidrug response in T-ALL cell lines

To determine a hPTM signature indicative of epidrug sensitivity in T-ALL, we integrated drug response profiles of 21 T-ALL cell lines towards nine epidrugs with the baseline global hPTM levels of the cell lines, as derived from our previously published MS-based atlas^15^. **Figure 1** presents a heatmap illustrating the summarized z-scores of the abundances of individual hPTMs quantified using msqrob2PTM^16^ across 21 T-ALL cell lines, colour-coded by their genetic subtypes. The modifications analyzed include acetylation (Ac), methylation (Me2, Me3), butyrylation (Bu), and phosphorylation (Ph), measured at specific lysine (K) or arginine (R) residues. Note that butyrylation is induced during sample preparation through propionylation of monomethylated lysines and is therefore interpreted as the latter. The heatmap reveals substantial heterogeneity in hPTM profiles among the T-ALL cell lines, with no evident clustering or correlation with their genetic subtypes. Yet, we have previously shown that the hPTM signature of T-ALL cell lines remains remarkably stable between two independent experiments measured 2 years apart, underscoring their reproducibility^15^. These findings suggest that hPTMs may act as independent markers of phenotype and therefore potentially therapeutic response, driven more by the cellular environment than by genetic subtype.

**Figure 1.**
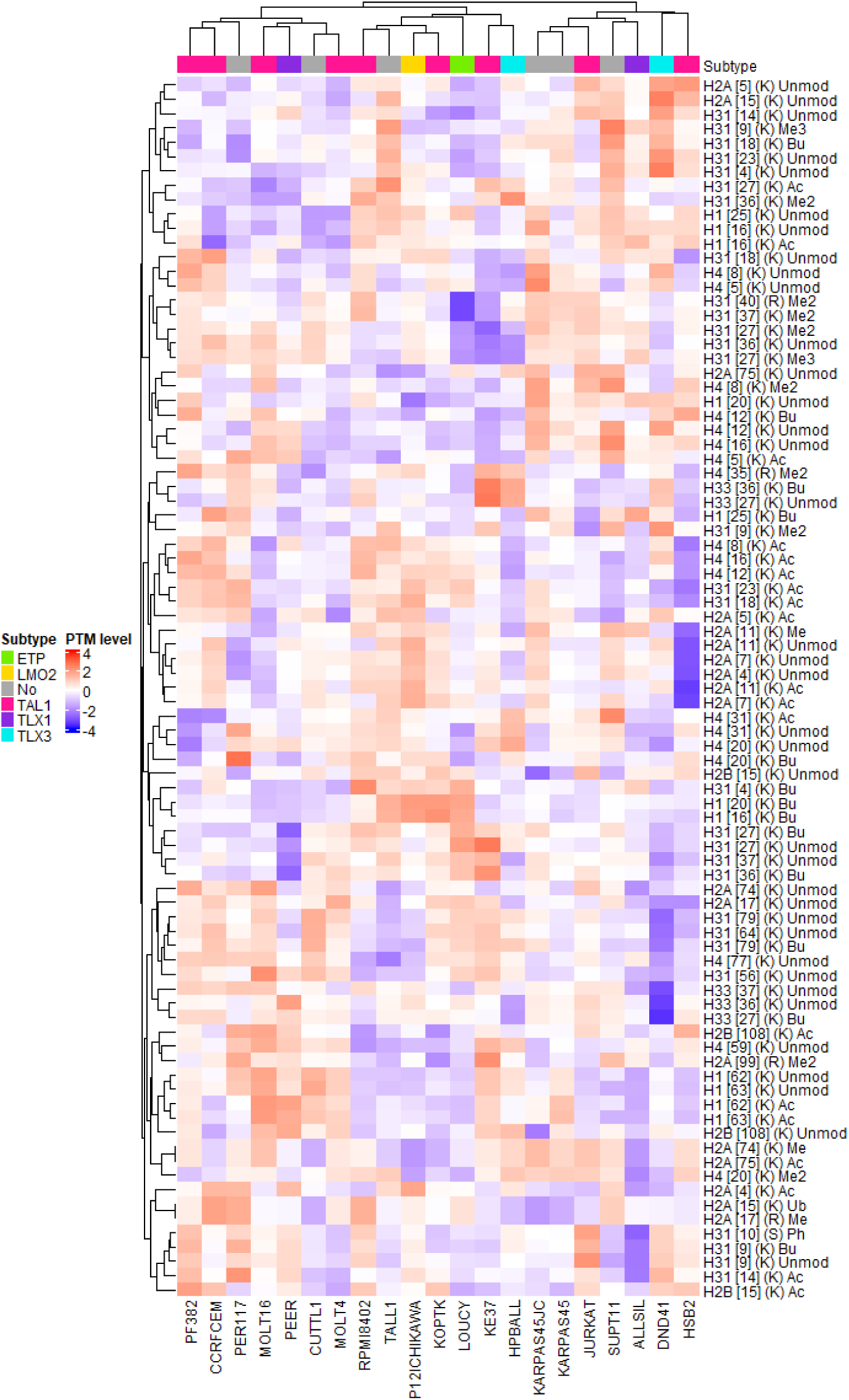
Clustered heatmap showing normalized and summarized single PTM levels for 21 T-ALL cell lines.

To generate drug response profiles, the 21 T-ALL cell lines were treated *in vitro* with a dilution series of nine epidrugs subdivided in three drug classes relevant to T-ALL therapy, i.e. anthracyclines, histone deacetylase (HDAC) inhibitors and DNA methyltransferase (DNMT) inhibitors. The anthracyclines doxorubicin (DOXO) and daunorubicin (DAUNO) are commonly used in the clinic as first-line treatments for T-ALL patients, while aclarubicin has also shown promising anti-leukemic effects in several preclinical studies^17–19^. Anthracyclines were thought to act primarily through DNA intercalation and inhibition of topoisomerase enzymes. However, it was recently discovered that anthracyclines can evict histones from nucleosomes^20,21^. ACLA mainly evicts histones from H3K27me3-labeled chromatin, whereas DOXO and DAUNO preferentially evict histones from H3K36me3-labeled chromatin^22^. In contrast to DOXO and DAUNO, ACLA primarily induces chromatin damage without causing double strand breaks, thereby avoiding cardiotoxicity while maintaining anticancer potency. The HDAC inhibitors used, namely Panobinostat (PAN), vorinostat (VOR) and romidepsin (ROM), have been shown to successfully target altered acetylation and increased levels of HDACs in hematological cancers such as T-ALL^23–29^. By influencing the acetylation level on histones, transcription factors or their essential co-factors, HDAC inhibitors have an indirect effect on gene expression. The third class of tested epidrugs, DNMT inhibitors, can be subdivided into nucleoside-analogues, including azacytidine (AZA) and decitabine (DAC), and non-nucleosides-analogues, such as the newly discovered drug GSK3685032 (GSK). Nucleoside analogues are non-selective inhibitors that target DNMT1, DNMT3A, and DNMT3B through irreversible covalent degradation of DNMT1, leading to DNA damage. In contrast, GSK3685032 acts as a selective inhibitor of DNMT1, operating through a reversible, non-covalent mechanism that does not cause DNA damage. AZA and DAC are currently used as a first-line therapy for the treatment of MDS, AML, JMML and CMML and have shown pre-clinical efficacy in T-ALL, whereas GSK3685032 recently showed improved tolerability and efficacy compared to the nucleoside-analogues in AML^30^. Although the primary mechanism of action of these DNMT inhibitors is to target the abnormal DNA methylation profile, AZA and DAC have recently been shown to also cause histone PTM changes^31^.

Based on the cell viability data, dose-response curves were generated for each drug (**Supplementary Figure 1**), and IC50 values were calculated (**Supplementary Table 3**). To identify associations between epigenetic marks and drug sensitivity, we employed two complementary approaches. First, a correlation-based analysis was performed by calculating Spearman correlations between hPTM levels and IC50 values across all 21 cell lines (**Figure 2A, Supplementary Table 4**). Herein, positive correlation coefficients indicate that high hPTM levels are associated with higher IC50 values, reflecting lower sensitivity to the drug. Conversely, negative correlation coefficients indicate that high hPTM levels correspond to lower IC50 values, linking these hPTMs to higher sensitivity. Since Spearman correlation considers only the ranks of hPTMs and IC50 values, differences in drug potency, such as between vorinostat and panobinostat, do not impact the analysis. Second, we conducted a differential hPTM analysis by comparing the top *n* most sensitive versus the least sensitive cell lines, thereby focusing on a subset of the data to highlight more pronounced differences (**Figure 2B, Supplementary Table 5**). While correlation analysis captures continuous associations across the entire dataset, it relies on sufficient variation in both variables and is sensitive to outliers. Conversely, differential analysis is more robust to noise and easier to interpret, but at the cost of reduced sample size and potential threshold bias. Applying both approaches in parallel enabled a more comprehensive assessment of hPTM–drug response relationships.

**Figure 2.**
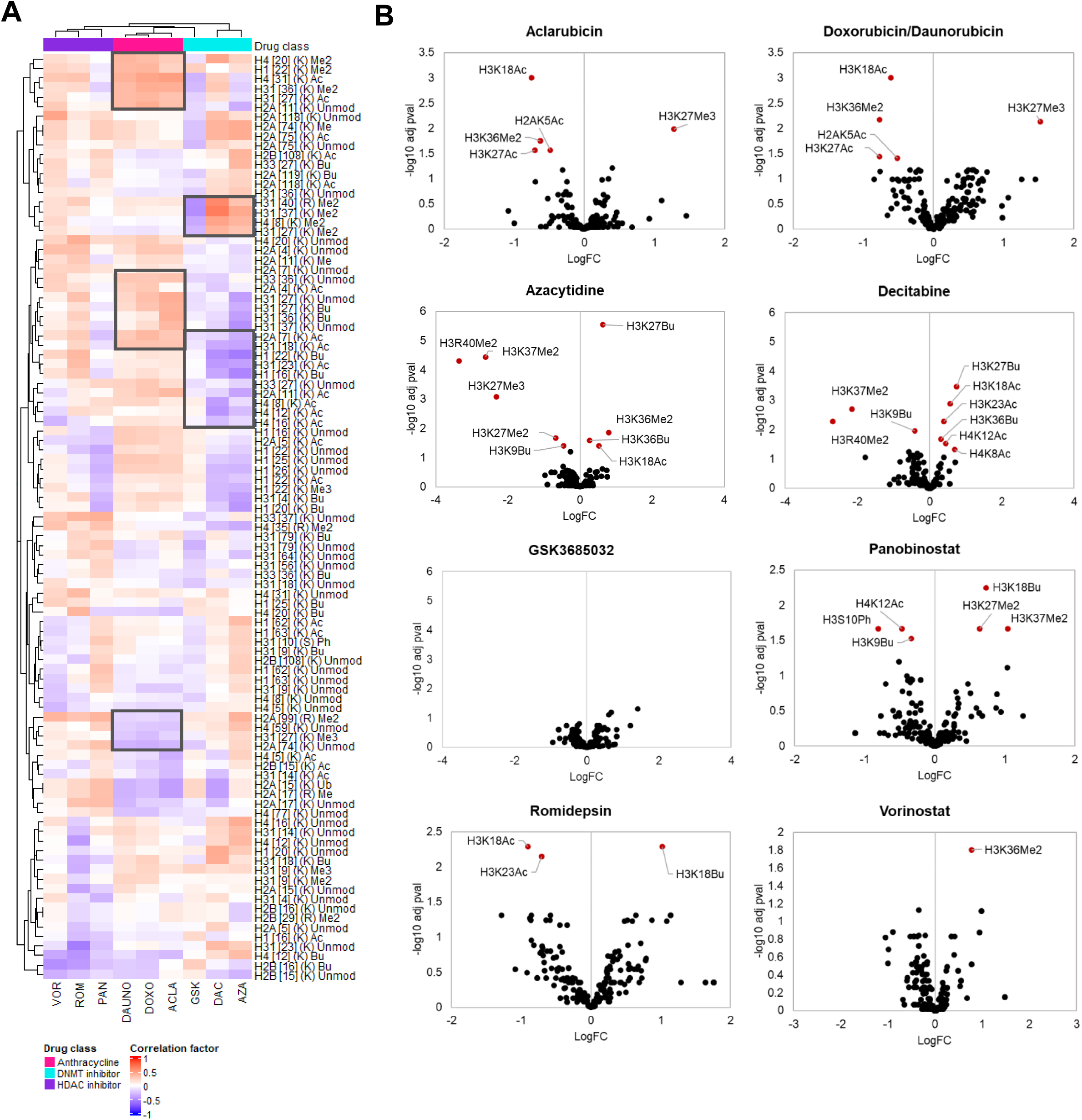
Epidrug sensitivity can be linked to hPTM levels in T-ALL cell lines. **(A)** Clustered heatmap showing the Spearman correlation factors between hPTMs (rows) and IC50 values (columns) over the 21 cell lines. Positive correlation coefficients (red) indicate that higher hPTM levels are associated with higher IC50 values, suggesting a link to drug resistance. Conversely, negative correlation coefficients (blue) imply that high hPTM levels are associated with lower IC50 values, indicating a potential association to drug sensitivity. **(B)** Volcano plots depicting the differential analysis between the least sensitive (left) and most sensitive (right) cell lines for each drug. Significantly enriched (adj pval < 0.05) PTMs are shown in red.

Hierarchical clustering of hPTM-IC50 Spearman correlation values revealed clear grouping of drugs within the same class, except for GSK (**Figure 2A**). This was particularly evident for the anthracyclines, which exhibited highly similar sensitivity patterns across individual compounds. This reflects their highly consistent IC50 rankings, with the same cell lines showing the highest sensitivity to all three anthracyclines (**Supplementary Figures 1–2**). As a result, the resolving power of the anthracycline class is effectively equivalent to that of a single compound, as their behavior is nearly indistinguishable, akin to triplicate measurements of the same drug. Despite the loss of resolving power within class, there was a clear distinction between more sensitive and less sensitive cell lines, reflected in a broader spectrum of IC50 values, which enabled the calculation of a more accurate Spearman correlation. For example, some cell lines, such as KOPT-K1, HSB-2, MOLT-16, and MOLT-4, were significantly more susceptible to anthracycline treatment compared to less sensitive lines like TALL-1. Briefly, H3K27Me3 was linked to higher sensitivity as previously reported, while mostly dimethylation marks such as H3K36Me2 and H4K20Me2 were linked to lower sensitivity (highlighted in **Figure 2A**). As DOXO and DAUNO share the same subset of top sensitive cell lines, they are shown together in the volcano plot. Interestingly, across all anthracyclines, the same hPTMs showed differential usage, with only one mark significantly enriched in the more sensitive cell lines, namely H3K27Me3 (adj. pval < 0.05). In contrast, acetylation at H3K18, H3K27 and H2AK5 were significantly enriched in the less sensitive cell lines, as well as H3K36Me2. The enrichment of H3K36me2 in this group is consistent with its known inverse relationship with H3K27me3^32^.

A similar pattern was observed for AZA and DAC, where overlapping sensitivity profiles across cell lines resulted in a correlation between their IC50 values (**Supplementary Figures 1–2**). In contrast, no such correlation was observed for GSK, likely due to its distinct and more selective mechanism of action, which targets a different subset of sensitive cell lines. For instance, KOPT-K1 and TALL-1 showed low IC50 values for both DAC and AZA, but high IC50 values for GSK, whereas KARPAS-45 exhibited the opposite pattern. Although DAC demonstrated lower efficacy than AZA, as it did not reduce cell viability to 0% in most cell lines, its potency was higher based on the lower IC50 values seen for DAC compared to AZA, at least in this in vitro experimental setup. Notably, treatment with GSK did not result in a reduction of cell viability to 0% for any of the cell lines, suggesting that GSK has reduced efficacy in T-ALL cell lines compared to DAC and AZA. Spearman analysis revealed positive correlations between dimethylation of histone H3 and H4 N-tails and IC50 values for AZA and DAC, indicating that a high histone dimethylation level can be linked with lower sensitivity to these DNMTis (highlighted in **Figure 2A**). Interestingly, these correlations were reversed for GSK, further underscoring the different mechanisms of action. In contrast, N-terminal acetylation marks on histones H3 and H4 were negatively correlated with IC50s for all DNMTis, reflecting an association with increased sensitivity, albeit less pronounced for GSK. For AZA and DAC, differential analysis showed a high overlap in significantly enriched hPTMs (adj. pval < 0.05), as shown in **Figure 2B**. Dimethylation of H3K37 and H3R40 was enriched in the least sensitive cell lines, whereas H3K27Bu was enriched in the most sensitive cell lines. Unfortunately, no significantly differential hPTMs were identified for GSK.

HDAC inhibitors showed limited resolution in sensitivity between the cell lines (**Supplementary Figure 1**), reducing the reliability of Spearman correlation calculations and limiting the predictive power of hPTM abundances for sensitivity to these drugs. Nevertheless, hPTM-IC50 correlations showed similar patterns across the different HDACis, resulting in their co-clustering in the heatmap, with VOR and ROM forming a distinct subcluster (**Figure 2A**). Given the narrow dynamic range in IC50 values, a group-wise comparison of the most and least sensitive cell lines offers a more suitable approach to detect meaningful hPTM differences here. Notably, as shown in **Figure 2B**, there is limited consistency in differential PTM patterns within the drug class. In cell lines with the highest IC50 for PAN, H4K12Ac was enriched, whereas H3K18Ac and H3K23Ac were enriched in those with the highest IC50 for ROM. Interestingly, H3K18Bu was enriched in sensitive cell lines for both PAN and ROM, suggesting a potential shared sensitivity-associated mark. For VOR, the only PTM associated with higher sensitivity was H3K36me2.

Of note, single hPTMs inferred using msqRob2PTM are derived from measurements of various histone peptidoforms, which may obscure combinatorial PTM patterns. However, similar conclusions were drawn when considering Spearman correlations between IC50 values and the identified peptidoforms (**Supplementary Figure 3, Supplementary Table 6**). In conclusion, we here show that hPTM signatures are associated with epidrug response in T-ALL cell lines, albeit without inferring causal relationships.

### Aclarubicin sensitivity is not solely determined by H3K27me3 levels

Our analysis identified H3K27me3 as the only significantly enriched hPTM in aclarubicin sensitive cell lines. This aligns with aclarubicin’s previously described preference for evicting histones from H3K27me3-marked heterochromatin, as well as the fact that aclarubicin was most effective in inducing apoptosis in DLBCL cells with high levels of H3K27me3^33^. To further investigate the relationship between H3K27me3 levels and aclarubicin sensitivity, we conducted cell viability assays following pharmacological inhibition of EZH2, the enzyme responsible for H3K27me3 deposition. Although pre-treatment with the EZH2 inhibitor GSK126 effectively reduced H3K27me3 levels (**Figure 3A, Supplementary Figure 4**), there was no observed change in the sensitivity of the cell lines to aclarubicin (**Figure 3B**). Furthermore, we identified T-ALL cell lines, such as JURKAT and KARPAS-45, which exhibited high H3K27me3 levels but low sensitivity to aclarubicin (**Figure 3A**). These results suggest that aclarubicin sensitivity in T-ALL cell lines is not solely dependent on H3K27me3 levels. Consequently, our findings underscore the importance of considering the broader histone modification landscape, rather than focusing on a single hPTM, when identifying biomarkers for drug sensitivity.

**Figure 3.**
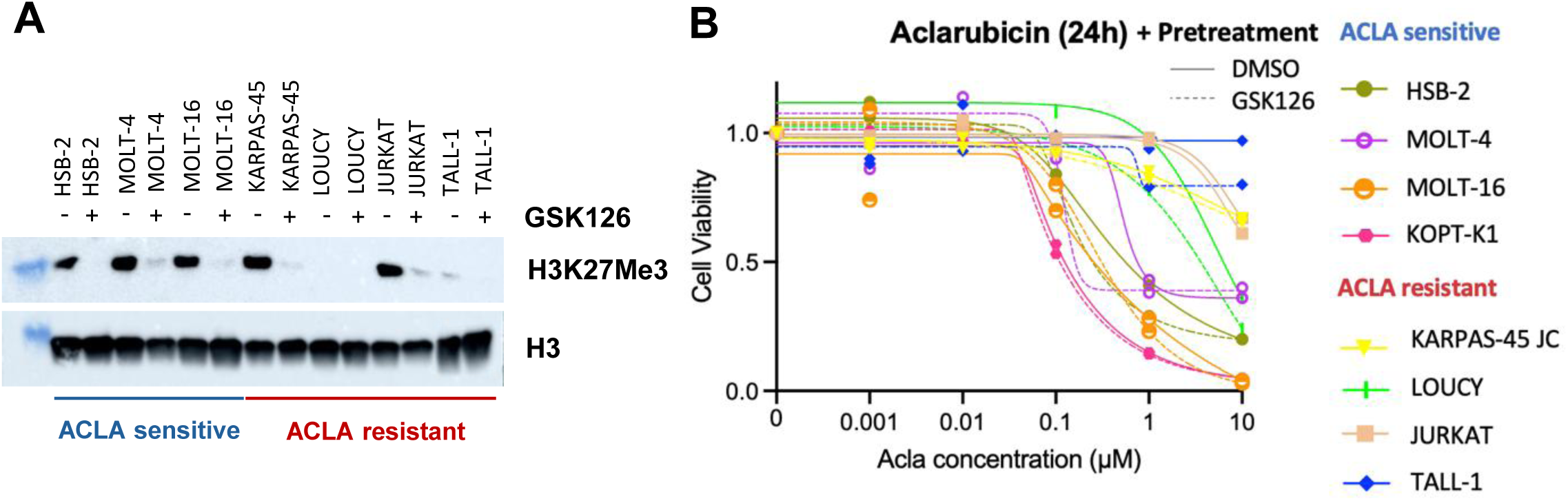
Inhibition of H3K27me3 does not alter aclarubicin sensitivity in T-ALL cell lines. **(A)** Western blot analysis visualizing H3K27me3 levels after 24-hour aclarubicin treatment in a subset of eight cell lines with or without 72-hour pre-treatment with the EZH2 inhibitor GSK126. Although the aclarubicin-sensitive cell line KOPT-K1 could not be included due to failed histone extraction, the results show a marked reduction in H3K27me3 levels in other cell lines after GSK126 treatment. Both the aclarubicin-resistant cell lines JURKAT and KARPAS-45 show high H3K27me3 levels despite resistance. **(B)** Cell viability curves for the four most sensitive and four least sensitive T-ALL cell lines in response to a dilution series of aclarubicin, with and without pre-treatment with GSK126. Each data point represents the mean of three independent experiments (see Supplementary Figure 4). The removal of H3K27me3 had no impact on the sensitivity of the cell lines to aclarubicin.

### Baseline hPTM profiling of T-ALL PDX models reveals epigenetic subclusters concordant with genetic subtype

To facilitate the clinical translation of hPTM signatures, it is essential to validate our findings in patient samples. However, primary T-ALL material is very sparse, often resulting in low-input quantities and thus requiring highly sensitive MS instruments for untargeted hPTM profiling. Previous epigenetic studies utilizing ATAC-seq and DNA methylation data have demonstrated that patient-derived xenografts (PDXs) serve as a reliable proxy for primary patient material in T-ALL, with correlations exceeding 0.9^34^. Therefore, we generated an hPTM atlas of 10 T-ALL PDX models (**Figure 4A**). In addition to hPTM profiling, we assessed global DNA methylation and identified the CpG Island Methylator Phenotype (CIMP) of the PDXs, which has previously been shown to have prognostic relevance in T-ALL^35,36^. Hierarchical clustering analysis of hPTMs revealed four main subclusters of T-ALL PDX models, partially driven by differential methylation of the histone H3 N-terminal tail, including the repressive marks H3K27Me3 and H3K9Me3 (highlighted in **Figure 4B**). Interestingly, this clustering pattern closely corresponded with the genetic subtype, and to some extent with CIMP status.

**Figure 4.**
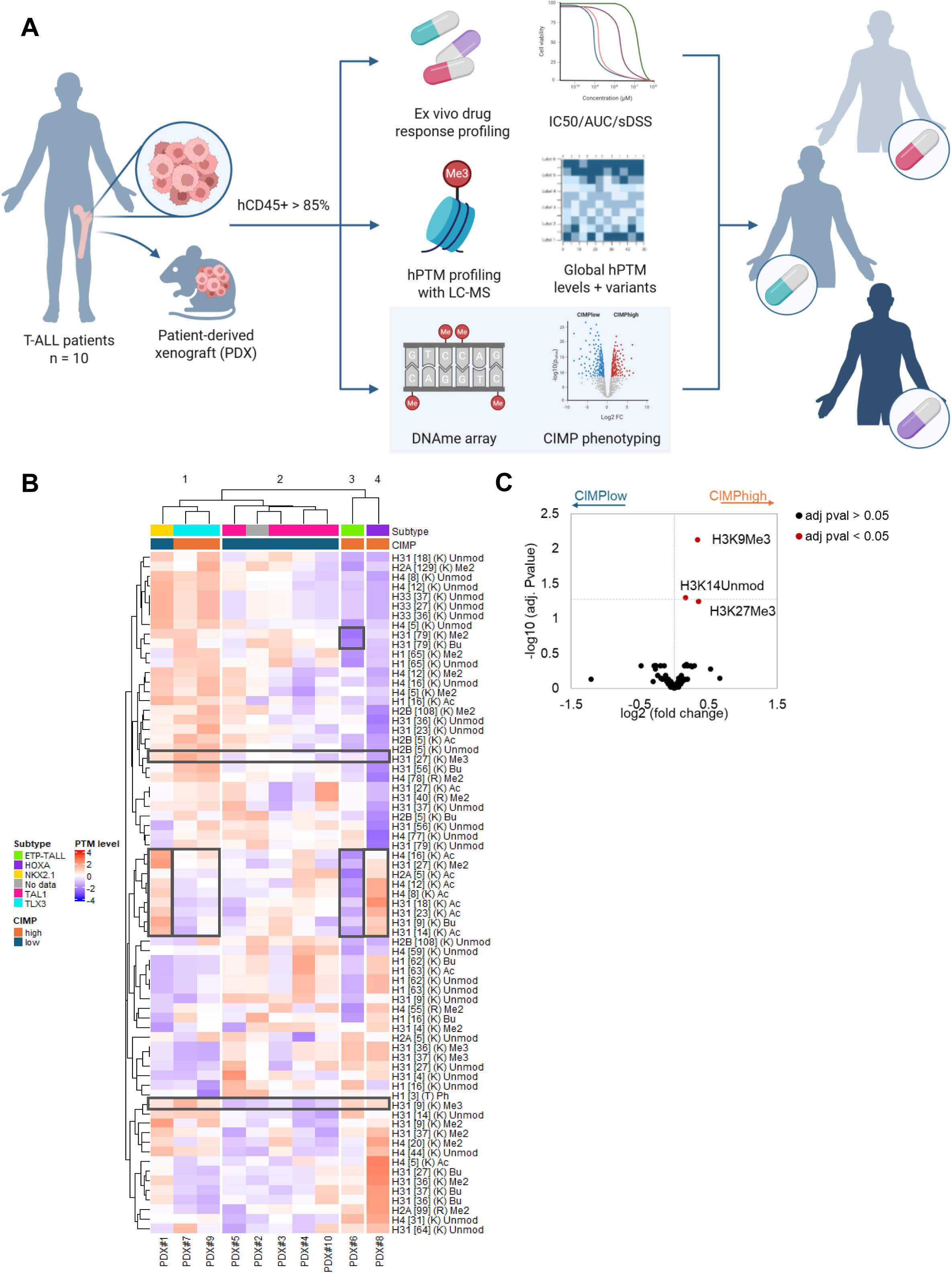
Epigenetic profiling of T-ALL PDX models. **(A)** NSG mice were engrafted with malignant primary cells derived from the bone marrow of 10 T-ALL patients for tumor expansion. Upon disease establishment, mice were sacrificed, and leukemic cells were harvested from the spleen. To mimic biological replicates, six mice were injected per patient. The percentage of human CD45+ cells (hCD45%) was measured by flow cytometry, with only samples exceeding 85% used for downstream analysis. Global hPTM profiling and DNA methylation sequencing were performed for all 10 PDX models. In parallel, PDX cells were treated ex vivo with a panel of eight epidrugs to assess drug sensitivity. Spearman correlation analysis was then conducted to explore the relationship between hPTM levels and drug response. Figure created with Biorender.com. **(B)** Clustered and annotated heatmap displaying the normalized and summarized single hPTM levels across the 10 PDX models. **(C)** Volcano plot illustrating the differential analysis of hPTMs between CIMPlow and CIMPhigh PDX models.

Cluster#1, consisting of two TLX3 T-ALLs and one NKX2.1 T-ALL, was characterized by high levels of the heterochromatin marks H3K9Me3 and H3K27Me3. The main difference between the two different subtypes within this cluster involved H3 and H4 acetylations (highlighted). Cluster#2 nicely corresponded with TAL1 subtype and CIMPlow status and was mainly characterized by a low H3K9Me3 level. Cluster #3, associated with the early T-cell precursor (ETP)-ALL subtype, exhibited an overall remarkably distinct hPTM landscape compared to the others. ETP-ALL is a subtype that arises from very early T-cell precursors, characterized by an epigenetic landscape and transcriptional programs that are more similar to hematopoietic stem cells. More specifically, this cluster was characterized by notably low levels of H3K79 mono- and dimethylation, as well as reduced overall acetylation (highlighted). Finally, Cluster #4, which includes a HOXA-driven T-ALL case, closely resembles Cluster #3 but shows notably higher levels of H3 and H4 acetylation marks, along with some increased di- and trimethylation marks. HOXA genes, typically involved in embryonic development and hematopoietic stem cell self-renewal, are aberrantly activated in this subtype. Interestingly, HOXA-driven T-ALL frequently exhibits features of the ETP-ALL subtype^37^.

To complement the annotated heatmap, we performed a differential analysis of hPTMs between CIMPlow and CIMPhigh, indicating that H3K9Me3, H3K27Me3 and H3K14Unmod were significantly enriched in CIMPhigh PDX models (**Figure 4C**).

### hPTM sensitivity signature is dependent on disease model

Ex vivo treatment of the PDX models was performed using the same drugs as those applied to the cell lines, with the exception of GSK3685032, which was excluded due to its prolonged incubation time, making it unsuitable for primary patient-derived material. IC50 values were subsequently determined, which showed overall lower variability among the PDX models compared to cell lines (**Supplementary Table 7**, **Supplementary Figures 5-6**). While the smaller sample size for PDX models may influence correlation estimates, several notable shifts in compound response are observed compared to the cell culture data. Most strikingly, the previously coherent compound IC50 correlation within the anthracyclines is disrupted, with only DOXO and DAUNO maintaining a similar drug response profile in PDX models. Additionally, PAN and VOR exhibited increased similarity at the cost of the other HDACi binary correlations that were still significant in the cell line experiment.

To assess whether our in vitro observations were reflected in vivo, we compared two PDX models differing in H3K27me3 status; one high and one low, and observed in vivo sensitivity using doxorubicin as a representative example. Doxorubicin treatment was initiated when at least 20% of leukocytes were hCD45-positive. Blood samples were collected on days 1, 5, and 8 (the day of sacrifice) to measure the proportion of hCD45+ cells. Mice were sacrificed one week after treatment initiation, and the proportion of hCD45+ leukemic blasts was quantified using flow cytometry and compared with untreated controls to assess the treatment effect. Additional indicators of treatment response included the comparison of spleen weight and size between treated and control mice. Consistent with the ex vivo results, PDX1 exhibited greater sensitivity to doxorubicin than PDX2 (**Supplementary Figures 5-6**), supporting the use of this ex vivo approach for predicting drug sensitivity in PDX models.

Similar to the cell line analysis, Spearman correlations between IC50 values and hPTM levels were calculated (**Supplementary Table 8**, **Figure 5A)**. Interestingly, clustering analysis revealed two main clusters; DNMTis were split with AZA grouping with the anthracyclines, while DAC clustered with the HDACis. Despite yielding higher absolute correlation coefficients than in cell lines (potentially due to the lower number of samples), the previously identified hPTM signatures could not be validated. Moreover, certain hPTMs demonstrated inverse correlations in comparison to the cell lines. For example, H3K27Me3, previously identified as a sensitivity marker for aclarubicin in cell lines, now exhibits an opposite trend in PDX models, with even sensitive PDX models displaying low levels of this modification. **Figure 5B** presents volcano plots showing a complementary differential analysis between the most sensitive and least sensitive PDX models for each drug (**Supplementary Table 9**). PAN and VOR are displayed together, as they share the same subset of PDX models. While not statistically significant, the top-ranking PTMs for each drug are labeled to highlight potential trends. Among all comparisons, H3K27me3 emerged as the only significantly differential PTM, showing enrichment in the least sensitive cell lines for AZA. Interestingly, H3K27me3 was also enriched in the least sensitive cell lines for ACLA, consistent with the correlation analysis, where these drugs clustered together. In contrast, the top-ranked PTMs for other drugs were not commonly shared within the same drug class, suggesting limited class-wide PTM signatures. Again, similar conclusions were drawn when calculating Spearman correlations between IC50 values and histone peptidoforms (**Supplementary Figure 8, Supplementary Table 10**).

**Figure 5.**
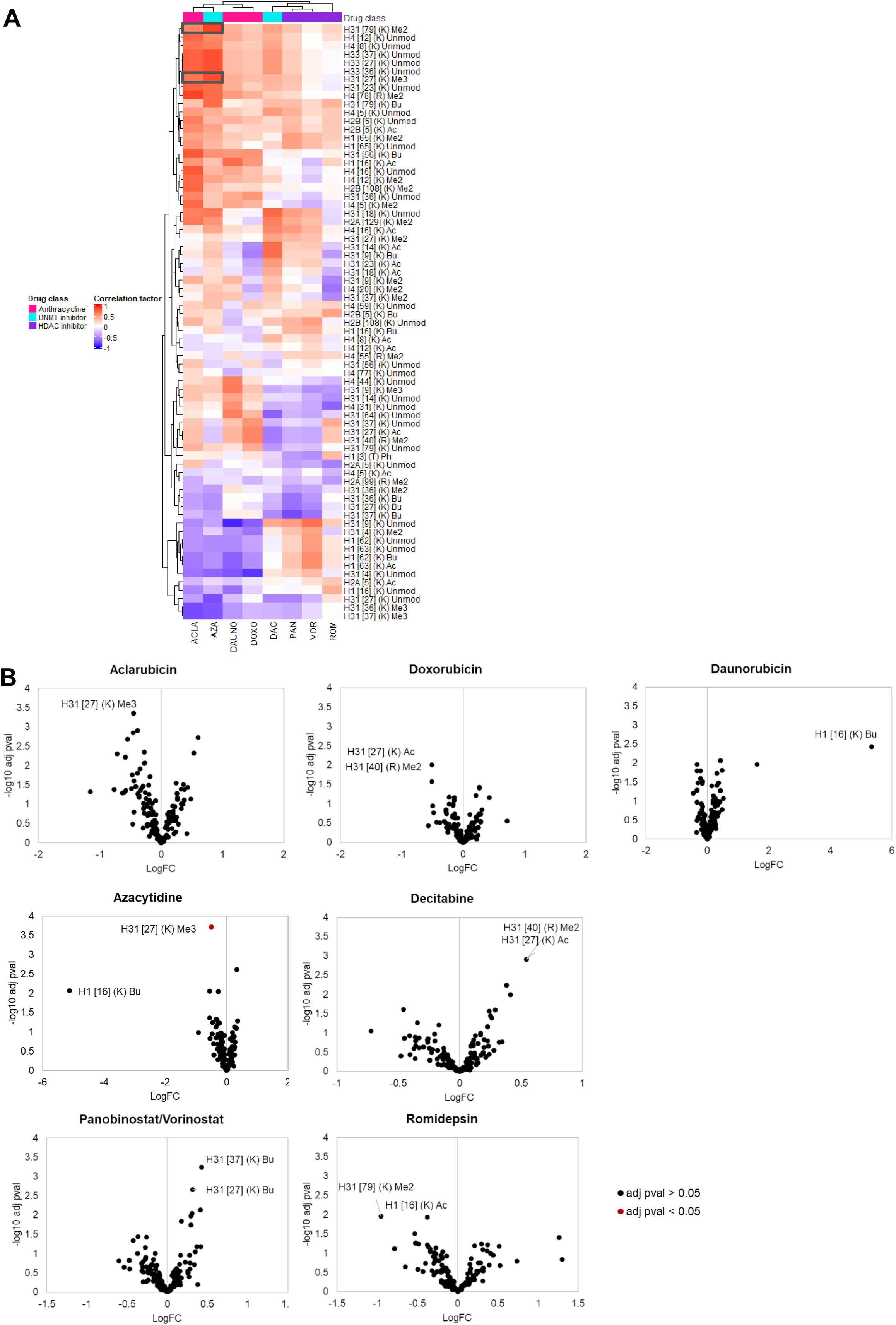
Ex vivo treatment of T-ALL PDX models. **(A)** Clustered heatmap showing the Spearman correlation factors between hPTMs (rows) and IC50 values (columns) over the 10 T-ALL PDX models. **(B)** Volcano plots depicting the differential analysis between the most resistant (less sensitive) (left) and most sensitive (right) PDX models for each drug. Significantly enriched (adj pval < 0.05) PTMs are shown in red.

Although it is well-recognized that cell lines do not fully replicate in vivo conditions, they remain a prevalent model in fundamental biological research. However, our data suggest that cell lines may not serve as a reliable proxy for predicting drug sensitivity based on hPTMs.

### Correlation analysis of hPTMs reveals divergent covariation in PDX models and cell lines

In recent years, the concept of a fixed “histone code” has been increasingly challenged, as individual hPTMs do not produce uniform outcomes across contexts. Instead, their functional impact is influenced by surrounding modifications and chromatin context. Therefore, analyzing hPTM covariation can serve as a proxy for their contextual interpretation in the cell and may help explain the discrepancies between cell lines and PDX models observed in our data. To explore this further, we quantified the pairwise Pearson correlation among common hPTMs across cell lines and PDX models to identify patterns of co-regulation. As shown in the correlation matrices in **Figures 6A-B**, the inverse correlating patterns are somewhat overlapping to the left of the correlograms, yet PDX models demonstrate a higher incidence of positively correlating hPTMs compared to cell lines, suggesting that the underlying epigenetic landscapes in these models differ significantly. H31K18Ac is highlighted to facilitate manual inspection. To further aid interpretation, we visualized hPTM co-variation using network graphs, where nodes represent individual hPTMs and edges represent significant Pearson correlations **(Figures 6C–D).** This approach revealed that acetylation marks (highlighted in purple) formed more tightly interconnected clusters in PDX models compared to cell lines, suggesting enhanced co-regulation. Additionally, PTMs on the H3.3 histone variant exhibited stronger correlations with other modifications in PDX models, pointing to broader integration into the epigenetic network. Notably, H3K79Bu emerged as a central hub (high-degree node) in cell lines, potentially indicating a key regulatory role in this context, whereas its centrality was reduced in PDX models. Conversely, H2AR99Me2 appeared as a major hub in PDX models, despite being a relatively understudied modification, suggesting a potentially unrecognized regulatory role in these systems.

**Figure 6.**
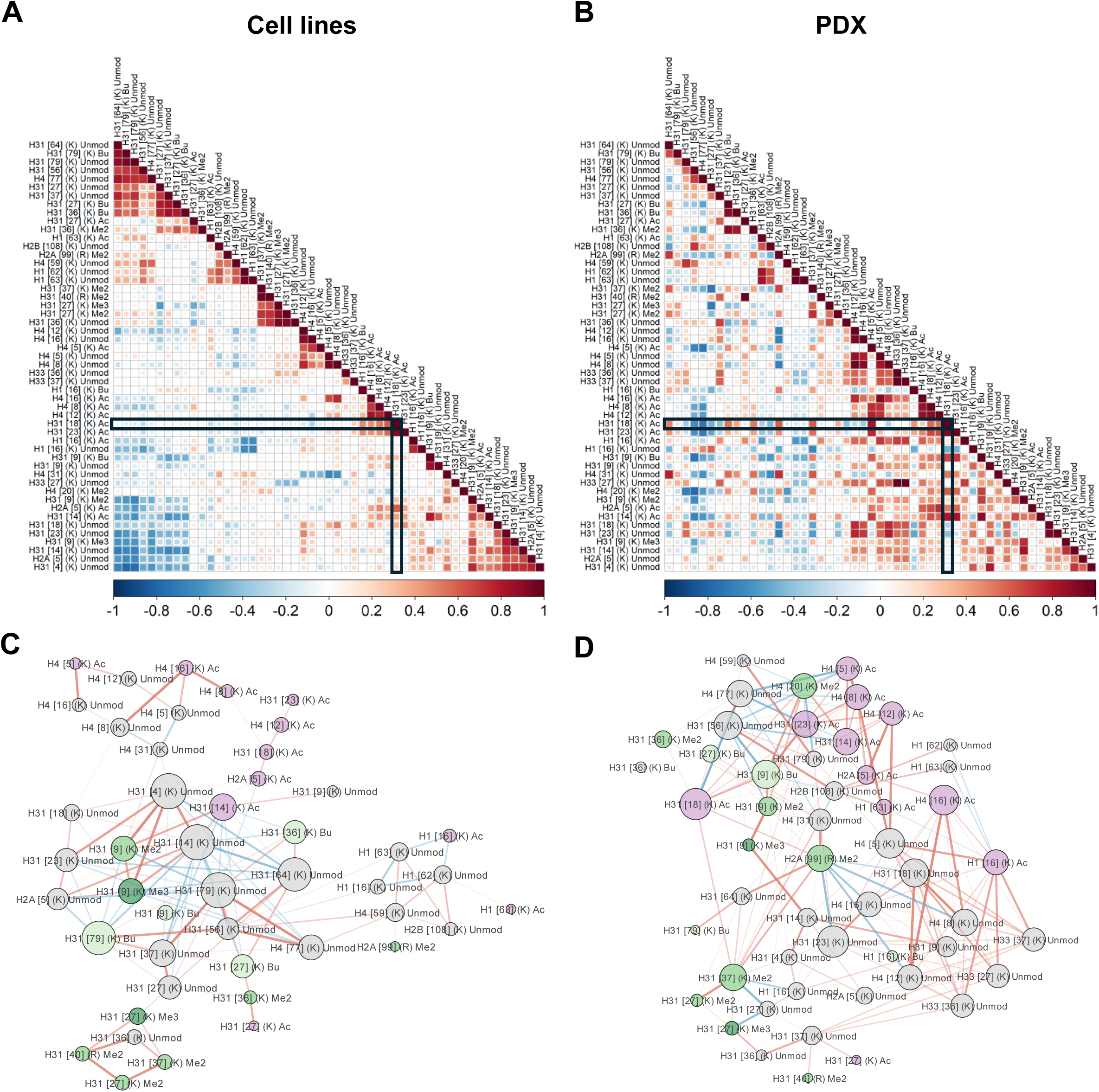
Correlation analysis of hPTMs reveals divergent covariation in PDX models and cell lines. **(A-B)** Covariance matrix of hPTM-hPTM Pearson correlations for cell lines **(A)** and PDX models **(B).** Network diagram derived from the covariance matrix in (A), showing significant Pearson correlations (p-value < 0.05, correlation coefficient > |0.5|) between histone PTMs. Nodes represent individual hPTMs, with node size indicating the degree (number of significant correlations) and node color corresponding to PTM type. Edges represent correlations between hPTMs, where edge color indicates either positive (red) or negative correlation (blue), and edge width reflects the absolute correlation coefficient.

## Discussion

Through a diverse array of chemical modifications, hPTMs dynamically shape the epigenetic landscape and influence transcriptional programs within tumor cells. Due to their pivotal role in gene regulation and their reversible nature, hPTMs have emerged as promising drug targets in cancer therapy ^38–40^. However, the combinatorial complexity of the hPTM code has hindered a comprehensive understanding of histone epigenetic dynamics in cancer, contributing to the inconsistent efficacy of epidrugs in clinical treatment.

Here, we aimed to apply an untargeted approach to investigate the hPTM profiles of 21 different T-ALL cell lines measured through MS, with the goal of evaluating the potential of hPTM patterns in predicting reduced sensitivity to epidrugs. By directly correlating hPTM abundances with the IC50 values of nine different compounds across three drug classes, we gained insights into several hPTMs of potential interest. However, the strength of these correlations was heavily influenced by the variation and consistency within the data matrix. Nevertheless, many of the histone marks identified here have previously been associated with cancer biology or epidrug response and could serve as interesting targets for future mechanistic studies.

For example, Neefjes et al. demonstrated that aclarubicin induces histone eviction specifically from H3K27me3-marked chromatin, suggesting that H3K27me3 could serve as a biomarker for aclarubicin sensitivity^41,42^. In our cell line analysis, we observed a slightly positive correlation between H3K27me3 levels and aclarubicin sensitivity. Despite the observed correlation, inhibiting the H3K27me3 writer EZH2 did not result in significant changes in aclarubicin sensitivity, suggesting that other factors likely influence drug response. Moreover, we saw an opposite trend in PDX samples, where H3K27me3 was significantly enriched in the least sensitive PDX models. We hypothesize that drug sensitivity is mediated by a network of interacting hPTMs rather than being driven by a single modification. For DNMTis we would expect that cell lines with inherently low levels of heterochromatin-associated marks would be more susceptible to treatment, as reduced DNA methylation, and thus decreased heterochromatin formation, would be less effectively compensated in these cells. Consistent with this, heterochromatin marks such as H3K9me3 and H3K27me3 were modestly enriched in the least sensitive cell lines, indicating that a more repressive chromatin state may confer lower sensitivity to DNMT inhibition. Conversely, for HDACis, cell lines with already high levels of histone acetylation may be less responsive, as the potential for further increasing acetylation is limited, thereby reducing the relative impact of the treatment. This pattern was evident in our data, with H4K12Ac enriched in the least sensitive cell lines for PAN, and H3K18Ac and H3K23Ac enriched in least sensitive lines for ROM, suggesting that pre-existing acetylation marks may diminish the cellular response to HDAC inhibition.

Rather than focusing solely on baseline hPTM levels, an alternative approach would be to examine how hPTMs change upon treatment in sensitive versus resistant cell lines, potentially revealing mechanistic insights or alternative explanations for drug response. For instance, in the case of decitabine, a previous MS-based study in two leukemia cell lines (MDS-L and TF-1) identified distinct treatment-induced changes in histone marks^43^. In MDS-L cells, H3.3K36me3 and the combinatorial acetylation H4K8acK12acK16ac were enriched in sensitive cells following decitabine treatment, whereas in TF-1 cells, increases in H3.1K27me1, H3.1K36me1, and the dual mark H3.1K27me1K36me1 were observed.

Cell lines have long been a cornerstone of drug development, but their relevance has been a subject of ongoing debate. While they retain critical cellular functions and fundamental molecular interactions, factors such as extended passaging, exposure to nutrient-rich media, and their immortalized nature can significantly alter molecular processes, diverging from their natural state at the time of isolation. To better approximate in vivo conditions, we chose to extend our analysis to PDX samples, including hPTM profiling, ex vivo IC50 measurements, and DNA methylation profiling. Due to the large number of compounds and models involved, conducting this analysis entirely in vivo was not feasible. However, to validate our ex vivo findings, we evaluated one compound in two in vivo PDX models. Interestingly, the PDX models maintained their sensitivity phenotypes ex vivo, as was previously shown in other studies ^44^.

Interestingly, based on their hPTM profiles, PDX samples clustered into four main groups, which showed clear alignment with genetic subtype, and to some extent CIMP signature. The link between CIMP and genetic subtype is in concordance with a previous study on a cohort of 109 T-ALL patients by Roels et al, which reported that CIMPlow T-ALLs are enriched for TAL1-rearranged cases, whereas CIMPhigh T-ALLs are enriched for HOXA, TLX3, and NKX2.1^45^. Our data largely mirror this pattern, with the exception of NKX2.1, which is represented by only one case in our study and just four cases in the Roels et al. cohort. Notably, the same study reported that CIMPlow T-ALLs exhibit high H3K27Me3 levels, whereas CIMPhigh T-ALLs show low H3K27Me3 levels. In contrast, we did not observe this pattern in our data. In fact, most CIMPlow cases displayed low H3K27Me3 levels, and two out of four CIMPhigh cases showed high H3K27Me3 levels. The differences between our findings and the previous study may be attributed to several factors. Our analysis was performed on only 10 PDX samples, while the study by Roels et al included 109 primary patient samples, which likely captures greater biological heterogeneity. Although they used flow cytometry and we used mass spectrometry to measure H3K27me3 levels, prior studies indicate that both methods provide comparable quantification, making methodological differences less likely to explain the variation^15^. On another note, the high level of H3K27Me3 observed in the TAL1 (CIMPlow) subcluster is in concordance with the selective epigenetic vulnerability to UTX/KMD6A demethylation of H3K27 of the TAL-1 subtype reported earlier^46^. Additionally, a distinct cluster was observed for both the ETP-ALL subtype (#3) and HOXA subtype (#4), both characterized by low H3K79Me2 and H3K27Me3 levels. ETP-ALL frequently harbors inactivating mutations in EZH2, the histone methyltransferase responsible for catalyzing H3K27Me3, consistent with its low level observed here. Loss-of-function mutations in *EZH2* increase transcription of stem-cell and early-progenitor related genes including *HOXA*, which in turn results into differentiation block of thymocytes^47,48^. A more targeted differential analysis of DNA methylation signatures, i.e. CIMPlow vs CIMPhigh PDX samples, revealed three hPTMs, H3K27me3, H3K14unmod, and H3K9me3, significantly enriched in the CIMPhigh group. This finding aligns with previous observations given the known association of both H3K9Me3 and DNA hypermethylation with heterochromatin ^45,49–51^. Similarly, H3K27me3, regulated by the Polycomb repressive complex 2 (PRC2), is crucial for gene repression in heterochromatic regions.

When correlating hPTMs with IC50 values for PDX samples, we observed that many of the hPTMs associated with reduced drug sensitivity in cell lines showed the opposite effect in PDX models. Notably, H3K27me3 inversely correlated with aclarubicin sensitivity in the ex vivo model. These findings suggest that T-ALL cell lines and PDX models may not be directly comparable in terms of hPTM–drug response relationships, or that specific hPTMs may be interpreted differently depending on cellular context. The functional impact of an hPTM is not binary but depends on its chromatin environment. Histone modifications function as part of a complex regulatory “language,” where their meaning is determined by interactions among writers, erasers, and readers within chromatin-modifying complexes. As a result, the same modification may correlate positively or negatively with phenotypic outcomes depending on the broader epigenetic landscape. This context-specific behaviour may explain the divergent hPTM–drug response patterns observed between cell lines and PDX models in our data. To investigate this further, we quantified the covariance between hPTMs across experimental conditions, essentially mapping hPTM coregulation. The results revealed that PDX samples exhibit a more tightly co-regulated histone code compared to the cell lines. In other words, a greater number of hPTM pairs demonstrated a high Pearson correlation across the PDX samples than in the cell lines. This “loosening of the histone network” in cell lines could reflect an adaptation to in vitro conditions, consistent with the changes in cellular processes that occur during immortalization and extended culturing ^52^. Moreover, a previous study that profiled and compared hPTMs using MS in patient tumor tissues, primary cultures, and cell lines from three different tumor models (breast cancer, glioblastoma, and ovarian cancer) revealed a significant rewiring of histone marks under cell culture conditions, with most changes being time-dependent and primarily observed in long-term cultures^53^.

In conclusion, while hPTMs hold great potential as biomarkers for predicting drug response, their utility is challenged by their inherent dynamic complexity and the significant influence of the chosen model system.

## Materials and methods

### Patient-derived xenograft generation

Lymphoprep-purified primary T-ALL cells from the blood or bone marrow of T-ALL patients were injected in the tail vein of immunodeficient NOD*.Cg-Prkdc^scid^ Il2rg^tm1Wjl^/SzJ* scid gamma (NSG, strain 005557, The Jackson Laboratory, ME, US) or *NOD.Cg-Prkdc^scid^ Il2rg^tm1Wjl^ Tg(CMV-IL3, CSF2, KITLG)1Eav/MloySzJ* (NSGS or NSG-SGM3, strain 013062, The Jackson Laboratory, ME, US) mice. To expand the model cells, primary PDX cells were further passaged by reinjecting leukemic cells in secondary NSG/NSGS mice. All human samples were acquired with written informed consent according to the Declaration of Helsinki, the studies were approved by the committee for medical ethics and laboratory animal experimentation of the Ghent University Hospital (UZGent; Ghent, Belgium) and Faculty of Medicine and Health Sciences of Ghent University (UGent; Ghent, Belgium). **Supplementary Table 1** summarizes the clinical and molecular metadata for the 10 patients samples used for xenograft generation.

### In vivo PDX treatment

Second generation PDX mice were used, starting from cryovials of the primary xenografts. For both PDX models, the same graft material was used for all experiments. Lymphoblast engraftment and leukemic burden was monitored by flow cytometric analysis on peripheral blood, using human CD45+ (hCD45) staining (CD45-FITC antibody; Miltenyi Biotec, Bergish Gladbag, Germany). To correct for lymphoid depletion induced by the treatment, Precision Count Beads (BioLegend, San Diego, CA, USA) were added to a fixed volume of peripheral blood. An exact number of hCD45+ cells per volume of peripheral blood (30 µL) was calculated using:

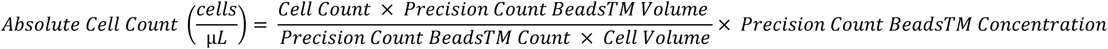

Once mice reached >20% hCD45+ leukocytes in peripheral blood, mice were randomized in two groups (N=4 mice/group). On day 1 and day 4 after randomization, mice were treated with either doxorubicin (0.6 mg/kg bodyweight in DMSO, intraperitoneal injection) or vehicle alone (DMSO, at a volume equivalent to that used for the 0.6 mg/kg doxorubicin dose). On day 8, mice were sacrificed, effect of treatment was analyzed based on spleen size and % spleen /body weight. Lymphoblast were collected from the spleen. In vivo experiments were approved by the ethical committee of laboratory animal experimentation of the Faculty of Medicine and Health Sciences of Ghent University.

### Cell culture

T-ALL cell lines (DSMZ, Braunschweig, Germany or gifted) were grown in culture flasks in RPMI 1640 medium (Gibco, Waltham, MA, USA) supplemented with 10% (HSB-2, JURKAT, LOUCY, RPMI-8402, DND-41, KOPT-K1, PER-117, PF-382, SUP-T11) or 20% (ALL-SIL, HPB-ALL, MOLT-16, PEER, TALL-1, CCRF-CEM, CUTTL1, KARPAS-45, KARPAS-45 JC, MOLT-4, KE-37, P12-ICHIKAWA) heat-inactivated fetal bovine serum (Sigma-Aldrich, St. Louis, MO, USA), penicillin (100 U/mL)-streptomycin (100 µg/mL) and 2 mM L-glutamine (Gibco, Waltham, MA, USA) and incubated at 37 °C with 5% CO2 and 95% humidity. Cultures were verified to be free of mycoplasma contamination using the TaKaRa PCR Mycoplasma Detection kit (TaKaRa Bio Europe, Goteborg, Sweden). **Supplementary Table 2** provides a comprehensive overview of the 21 cell lines, including their origin, immunophenotype, TCR status, translocations, oncogene classification and key gene mutations.

Patient-derived T-ALL cells from PDX mouse models were quickly thawed and stabilized for 24 hours before ex vivo treatment in U-bottom 96-well plates containing aMEM medium (Gibco, Waltham, MA, USA) supplemented with 10% heat-inactivated fetal calf serum, 10% heat inactivated human AB+ serum (Sigma-Aldrich, St. Louis, MO, USA), human IL7 (10ng/ml) (PeproTech®, Gibco, Waltham, MA, USA), human stem cell factor (50ng/ml) (PeproTech®, Gibco, Waltham, MA, USA), human FLT3 ligand (20ng/ml) (PeproTech®, Gibco, Waltham, MA, USA) and human IL2 (100ng/ml) (PeproTech®, Gibco, Waltham, MA, USA) and incubated at 37 °C with 5% CO2 and 95% humidity.

### Cell viability assays

T-ALL cells were plated in white 96-well plates at a density of 25,000 cells/well and treated with a six-point dilution series of azacytidine (AZA; Selleckchem, Houston, TX, USA), decitabine (DAC; Selleckchem, Houston, TX, USA), GSK3685032 (GSK; Selleckchem, Houston, TX, USA), panobinostat (PAN; Selleckchem, Houston, TX, USA), vorinostat (VOR; Selleckchem, Houston, TX, USA), romidepsin (ROM; Selleckchem, Houston, TX, USA), aclarubicin (ACLA; Lab of J. Neefjes, Leiden, The Netherlands), doxorubicin (DOXO; Selleckchem, Houston, TX, USA), and daunorubicin (DAUNO; Selleckchem, Houston, TX, USA). For AZA and DAC, each concentration was added daily during four days to assure constant drug exposure, since the half-life of AZA and DAC is approximately 8-12 hours in vitro. For each compound, the dilution series was optimized to center the concentration range around the predicted IC50. The treated cells were incubated for a duration between 1 and 6 days, depending on the epidrug (**Table 1**), at 37°C with 5% CO2 and 95% humidity. Next, cells were lysed with CellTiter-Glo (Promega, Madison, WI, USA) and luminescence was measured using a GloMax Discover Microplate Reader (Promega, Madison, WI, USA). Two technical replicates and three biological replicates were performed. For each cell line, the average luminescence and standard deviation of the three biological replicates were calculated at each concentration. Cell viability was calculated relative to vehicle-treated samples and plotted against compound concentration. Data were fit to generate a concentration response curve and absolute IC50 values were calculated for each compound using GraphPad Prism (GraphPad Software, San Diego, CA, USA).

**Table 1.** Overview of dilution series and incubation time per drug.

| Drug class | Epidrugs | Dilution series (in 2% DMSO – with exception of GSK in 4% ethanol) | Incubation time (days) |
| --- | --- | --- | --- |
| DNMT inhibitors | Azacitidine (AZA) | 50 – 10 – 1 – 0,1 – 0,01 – 0 (µM) | 4 |
|  | Decitabine (DAC) | 50 – 10 – 1 – 0,1 – 0,01 – 0 (µM) | 4 |
|  | GSK3685032 (GSK) | 10 – 5 – 1 – 0,5 – 0,1 – 0,01 – 0 (µM) | 6 |
| HDAC inhibitors | Panobinostat (PAN) | 25 – 10 – 5 – 2,5 – 1 – 0 (nM) | 3 |
|  | Vorinostat (VOR) | 5 – 1 – 0,75 – 0,5 – 0,25 – 0,1 – 0 (µM) | 3 |
|  | Romidepsin (ROM) | 5 – 2,5 – 1 – 0,5 – 0,1 – 0 (nM) | 3 |
| Anthracyclines | Aclarubicin (ACLA) | 10 – 1 – 0,1 – 0,01 – 0,001 – 0 (µM) | 1 |
|  | Doxorubicin (DOXO) | 10 – 1 – 0,1 – 0,01 – 0,001 – 0 (µM) | 1 |
|  | Daunorubicin (DAUNO) | 10 – 1 – 0,1 – 0,01 – 0,001 – 0 (µM) | 1 |

### H3K27Me3 inhibition experiment

A subset of the four most sensitive cell lines (HSB-2, KOPTK-1, MOLT-4, MOLT-16) and four least sensitive cell lines (JURKAT, LOUCY, TALL-1, KARPAS-45 (Lab of Jan Cools)) for aclarubicin were cultured as previously described and treated with either 1µM of GSK126 (Selleckchem, Houston, TX, USA) or vehicle (DMSO) for 72 hours. Following treatment, 2 x 10^6^ cells per condition were collected for western blot analysis. In parallel, a separate set of cells were seeded into 96 well plates (25,000 per well) and exposed to a dilution series of aclarubicin for an additional 24 hours. After treatment, cell viability was assessed using the CellTiter-Glo assay, as previously described.

### Western blot

Histones from 2 x 10^6^ cells were extracted via the Active Motif Histone Extraction Kit (Active Motif, Carlsbad, CA, USA). After extraction, histone concentration was measured via the Pierce BCA Protein Assay Kit (Thermo Fisher Scientific, Waltham, MA, USA). An equal amount of histones for each cell line were prepared for western blot. First, 5 µl 5x Laemmli buffer with β-mercaptoethanol (1/8) was added to 20 µl of sample and incubated for 10 minutes at 95°C for denaturation. Then, gel electrophoresis was performed by loading 10 µl of the samples and 4 µl of the Page Ruler Plus Prestained protein ladder (26620; Thermo Fisher Scientific, Waltham, MA, USA) on two 9-16% Mini-PROTEAN TGX Precast Protein Gels (Bio-Rad Laboratories, Hercules, CA, USA). To identify H3K27me3, the anti-H3K27me3 antibody (07-449; MilliporeSigma, Burlington, MA, USA) was added to one blot (1/1000 in 5% milk/TBST) and incubated overnight. A separate blot was probed with anti-H3 antibody (ab1791; Abcam, Cambridge, UK; 1:1000 in 5% milk/TBST) to provide a visual check of equal histone loading. Because H3K27me3 and total H3 were detected on separate blots, no per-lane normalization was performed between them. After addition of anti-rabbit HRP-linked antibody (1/10,000; Cell Signaling Technology, Danvers, MA, USA), both blots were visualized on the Amersham Imager 680 (GE healthcare, Chicago, IL, USA) via SuperSignal West Dura Extended Duration Substrate (Thermo Fischer Scientific, Waltham, MA, USA).

### DNA methylation array analysis

Genomic DNA was bisulfite-converted using the EZ DNA Methylation Kit or EZ DNA Methylation-Gold Kit (Zymo Research, CA, USA), following the manufacturer’s protocols. Methylation analysis was performed on Illumina platforms using Infinium MethylationEPIC v1.0 arrays (Illumina, CA, USA).

Raw methylation array data was pre-processed by setting CpG sites with bead coverage of ≤ 2 beads or a bead detection p-value of > 0.05 as missing. Bead exclusion was performed on non-CpG beads, beads within five base pairs of known European SNPs^54^, and multi-location probes^55^.

Data normalization was performed in two steps: first FunNorm from the R-package minfi^56^ was used for between-array normalization, followed by BMIQ from the ChAMP package^57^ for bead-type normalization. Beta values scaling from 0 (no methylation) to 1 (fully methylated) were computed. CpGs with missing values in > 50% of the samples were excluded while CpGs with missing data in < 50% of the samples had their beta values imputed using k-nearest neighbours imputation. Further filtering removed CpGs located on the X and Y chromosomes to minimize gender bias in downstream analysis. CpG Island Methylator Phenotype (CIMP) classification was performed using the previously established CIMP panel for EPIC v.1 arrays of 1,099 CpG sites^58^.

### Histone sample preparation

Histone extraction of the cells was performed using direct acid extraction as described previously^59^. Consequently, one-dimensional SDS-PAGE on a 9-18% TGX gel (Bio-Rad Laboratories, Hercules, CA, USA) was performed on a fraction of the resulting extracts for quantification and normalization^60^. The remaining histone extracts were propionylated and digested following an established protocol^61^.

### Liquid chromatography and mass spectrometry acquisition

The propionylated and digested samples were resuspended in 0.1% formic acid and injection volumes were adjusted resulting in 800 ng of histones on column. A quality control mixture was created by mixing 2 μl of each sample. Data-dependent acquisition (DDA) was performed on a ZenoTOF 7600 system (AB Sciex, Marlborough, MA, USA) operating in positive mode coupled to an ACQUITY UPLC M-Class System (Waters Corporation, Milford, MA, USA) operating in capillary flow mode (5 μl min*−1*). Trapping and separation of the peptides was carried out on a Triart C18 column (5 mm × 0.5 mm; YMC Europe GmbH, Dinslaken, Germany) and a Luna Omega Polar C18 column at 45°C (150 mm × 0.3 mm; particle size 3 µm; Phenomenex, Torrance, CA, USA), respectively, using a low pH reverse-phase gradient. Buffers A and B of the mobile phase consisted of 0.1% formic acid in water and in acetonitrile (Biosolve Chimie, Dieuze, France), respectively. A 20-min non-linear gradient going from 2% to 50% Buffer B, followed by a washing step at 90% mobile phase B and an equilibration step at 2% mobile phase B (starting conditions). The samples were run in a randomized fashion and a quality control injection was incorporated every ten samples. For each cycle, one full MS1 scan (m/z 350–1,250) of 100 ms was followed by an MS2 (m/z 140–1,800, high-sensitivity mode) of 12 ms. A maximum of 40 precursors (charge state +2 to +6) exceeding 1500 c.p.s. were monitored, followed by an exclusion for 3 s per cycle. A collision energy of 12 V and a cycle time of 800 ms was applied. Ion source parameters were set to 4.5 kV for the ion spray voltage, 35 psi for the curtain gas, 15 psi for nebulizer gas (ion source gas 1), 60 psi for heater gas (ion source gas 2) and 200 °C as source temperature.

### Mass spectrometry data analysis

Analysis of raw MS/MS data was performed as previously described^62^. Briefly, raw data from all runs were imported all runs were aligned in a single experiment in Progenesis QIP 4.2 (Nonlinear Dynamics - Waters, Newcastle upon Tyne, UK) for feature detection. Next, the 20 MS/MS spectra closest to the elution apex were selected for each precursor ion and merged into a single ∗.mgf file to search in Mascot (Matrix Science Ltd, London, UK). Mascot is a database search engine that uses a probabilistic scoring algorithm to define peptide-to-spectrum matches (PSM). Two types of searches were performed on this file: 1) a quality search to verify the presence of non-propionylated standards (ß-gal), and to assess the extent of underpropionylation; and 2) an error tolerant search to identify the proteins present in the sample, and to determine the best set of 9 hPTMs for further analysis. The Mascot search parameters used for both searches are shown in **Table 2**. A new FASTA-database was generated based on the results from the error tolerant search for further analysis. Next, the three MS/MS spectra closest to the elution apex were merged into a single *.mgf and exported to perform a biological search in Mascot (**Table 2**). The search results (∗.xml-format) were imported back into Progenesis QIP 4.2 to annotate the features from which they originated. Features that were annotated as histone peptidoforms were manually validated by an expert to resolve isobaric near-coelution (**Figure 2**). Consequently, runs were normalized against all histone peptides in order to assure a constant histone protein abundance, since we aim to quantify changes in hPTMs and not in the expression of histones themselves. Outliers were removed based on normalization factor (greater than 10 and less than 0,3) and PCA clustering of all histone peptidoforms. Finally, deconvoluted peptide ion data (*.csv format) from all histones were exported from Progenesis QIP 4.2 for PTM summarization using msqrob2PTM^16^.

**Table 2.** Overview of Mascot search parameters.

|  | Quality search | Error tolerant search | Biological search |
| --- | --- | --- | --- |
| Mass error precursor ions (ppm) | 10 | 10 | 10 |
| Mass error fragment ions (ppm) | 50 | 50 | 50 |
| Enzyme specificity | Trypsin | ArgC | ArgC |
| Miscleavages | 1 | 1 | 1 |
| Variable modifications | Propionylation (N-term)<br>Propionylation (K) | Deamidation (NQ)<br>Oxidation (M) | Acetylation (K)<br>Butyrylation (K)<br>Formylation (K)<br>Trimethylation (K)<br>Ubiquitination (K)<br>Methylation (R) |
|  |  |  | Dimethylation (KR)<br>Deamidation (NQR)<br>Phosphorylation (ST) |
| Fixed modifications | / | Propionylation (N-term)<br>Propionylation (K) | Propionylation (N-term)<br>Propionylation (K) |
| Database | Human Swissprot<br>supplemented with contaminants from the common Repository for Adventitious Proteins (cRAP) database | Human Swissprot<br>supplemented with contaminants from the common Repository for Adventitious Proteins (cRAP) database | Custom-made FASTA-database |

### Integration of drug response with hPTM data

A publicly available hPTM atlas comprising LC-MS data of 21 T-ALL cell lines was used to integrate with drug responses^15^. Briefly, raw abundances of all histone peptidoforms were exported from the Progenesis QIP project and summarized levels of single hPTMs were quantified using msqrob2PTM^16^. Differential analysis between the most sensitive and least sensitive cell lines was performed using linear regression models within msqrob2PTM. Spearman correlations between IC50s and PTM or peptidoform levels were calculated in R. Data visualisation, including heatmaps and volcano plots, was performed using custom R scripts.

### Covariation analysis

For both T-ALL cell lines and PDX models, a covariance matrix was calculated for all pairs of hPTMs to obtain Pearson R values of each PTM-PTM correlation and the significance of the correlations were estimated using a two-sided T-test. The lower triangle of each covariance matrix was visualized using the corrplot package^63^. Finally, significant correlations (p-value < 0.05) were visualized using Cytoscape^64^, with nodes representing hPTMs and edges representing correlation coefficients.

## Data and code availability

Cell line mass spectrometry raw data and analysis files were obtained from the ProteomeXchange Consortium (http://www.proteomexchange.org) under the dataset identifier PXD031500. Mass spectrometry raw data (*.wiff and *.scan files) and analysis files from the T-ALL PDX models generated in this study have been deposited to the ProteomeXchange Consortium via the PRIDE^65^ partner repository with the dataset identifier PXD067935 and 10.6019/PXD06793. The data is currently accessible with the token ywBECfsz6hWu and will be made publicly available after publication. All analysis scripts to process data and re-create figures are available at https://github.com/lcorvele/pharmacoepigenetics.

## Acknowledgement/Funding

This study has been supported by grants from The Research Foundation Flanders (FWO) awarded to L.C. (1SF2622N) and M.D. (12E9716N) and a preclinical research grant from Stand Up To Cancer (Kom Op Tegen Kanker-Flanders) awarded to P.V.V.

PDX models were generated with technical support from the PDXGhent core facility, with financial support from The Research Foundation Flanders (FWO) for medium-scale research infrastructure, Ghent University and Cancer Research Institute Ghent (CRIG).

Additional support was provided by The Swedish Childhood Cancer Foundation, the Swedish Cancer Society, the Cancer Research Foundation in Northern Sweden, and the Medical Faculty of Umeå University.

The PER-117 cell line was kindly provided by the PCH Oncology Biobank (Prof. Rishi S. Kotecha) which is funded by the Western Australian Future Health Research and Innovation Fund.

## Author contributions

L.C.: conceptualization, in vitro and ex vivo treatment, histone sample preparation, data acquisition, data analysis, writing. L.P.: conceptualization, in vitro treatment, PDX generation and in vivo treatment, writing. O.S.: in vitro treatment, writing. B.L.: cell culture. N.R. DNA methylation analysis. S.D.: DNA methylation analysis and co-supervision. M.L.: DNA methylation analysis. W.S.: PDX generation and biobanking. A.D.M.: data analysis and writing. B.D.M. and T.L.: collection of patient samples and clinical data collection. S.G.: co-supervision. D.D.: co-supervision. P.V.V.: conceptualization, supervision. M.D.: conceptualization, writing, supervision.

---

**Supplementary Figure 1.**
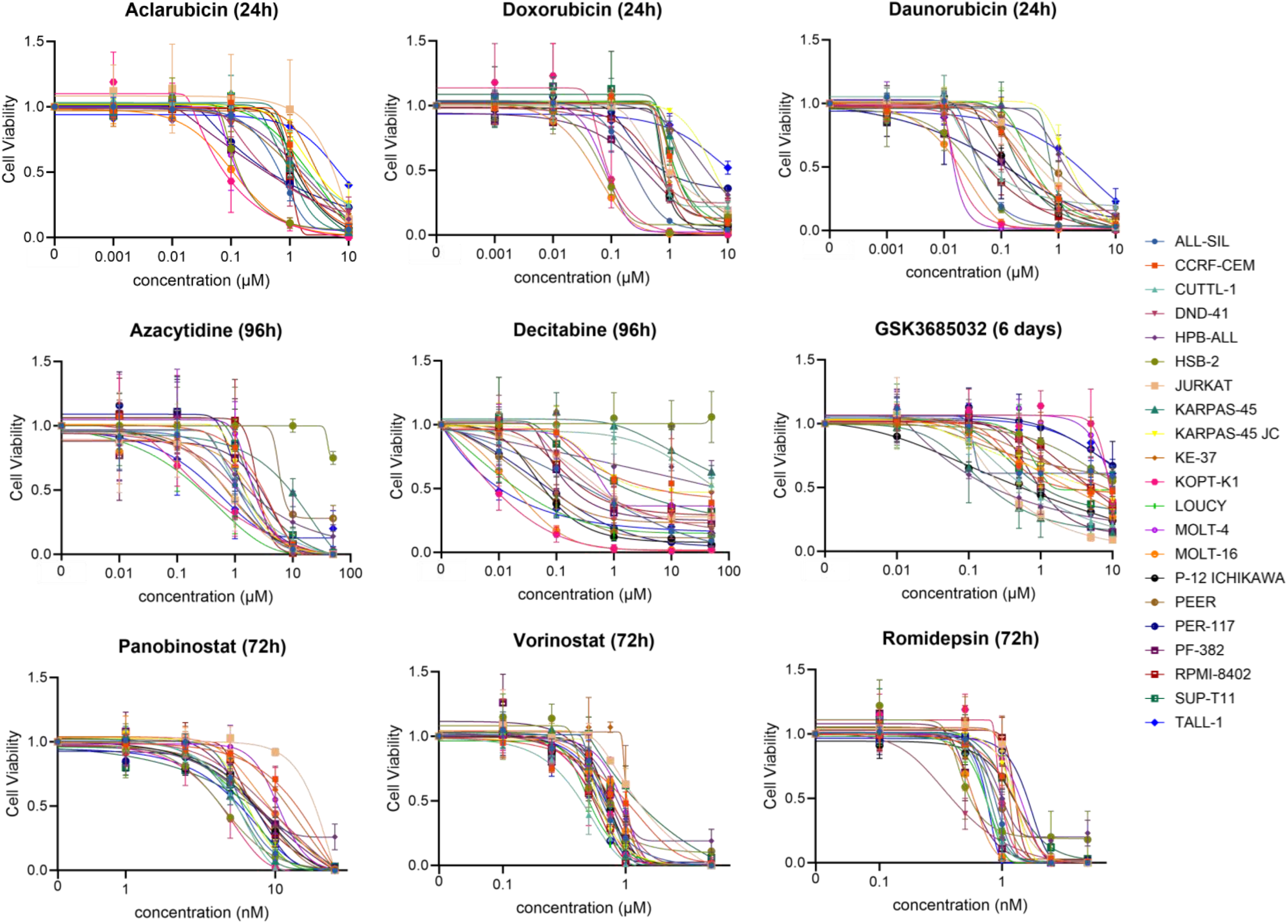
Dose-response curves showing the cell viability of 21 T-ALL cell lines (colors) after in vitro treatment with a dilution series of nine epidrugs. Each data point represents the mean of three independent experiments.

**Supplementary Figure 2.**
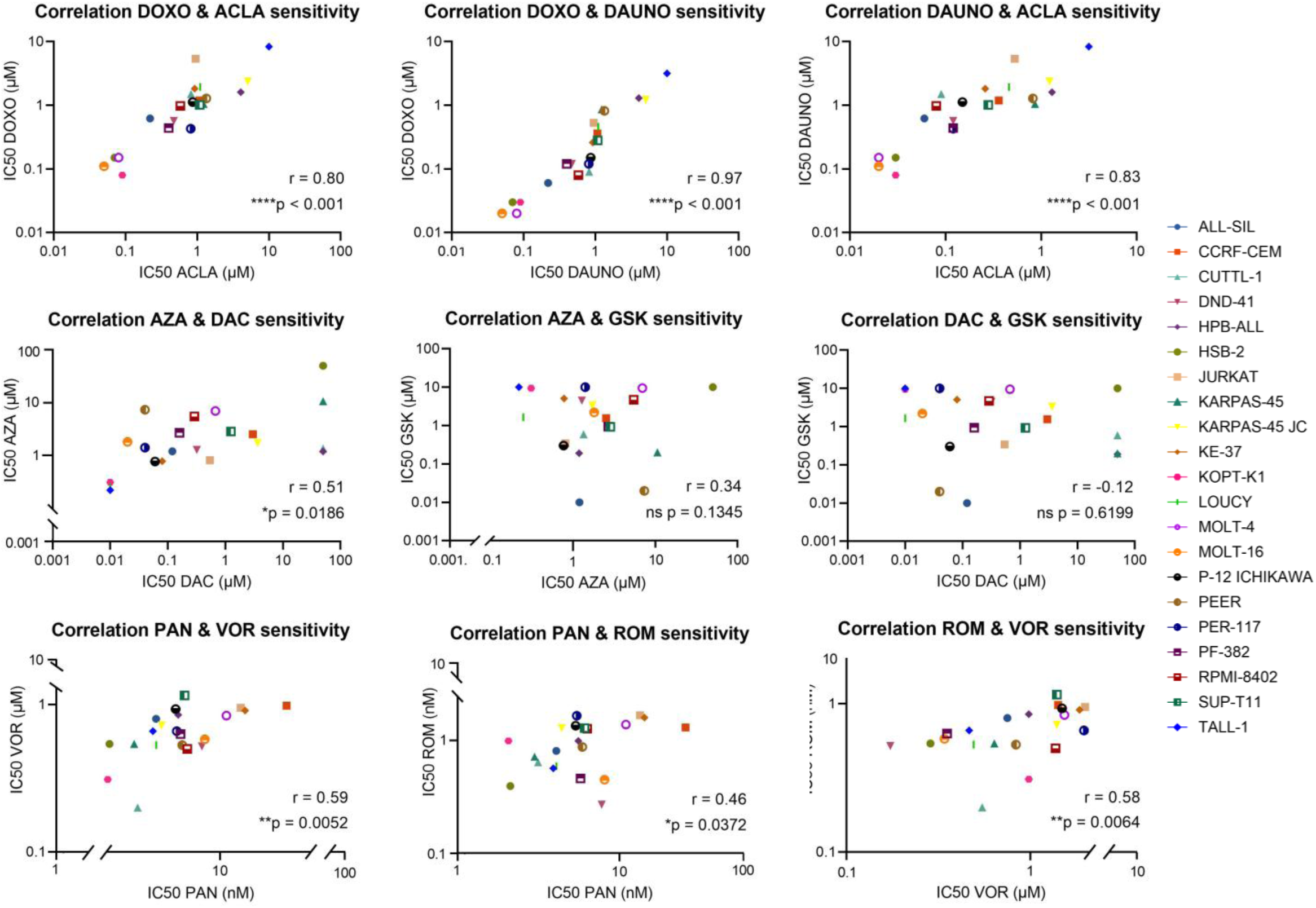
Scatter plots showing the Pearson correlation between the IC50s of 21 T-ALL cell lines (colors) within drug classes.

**Supplementary Figure 3.**
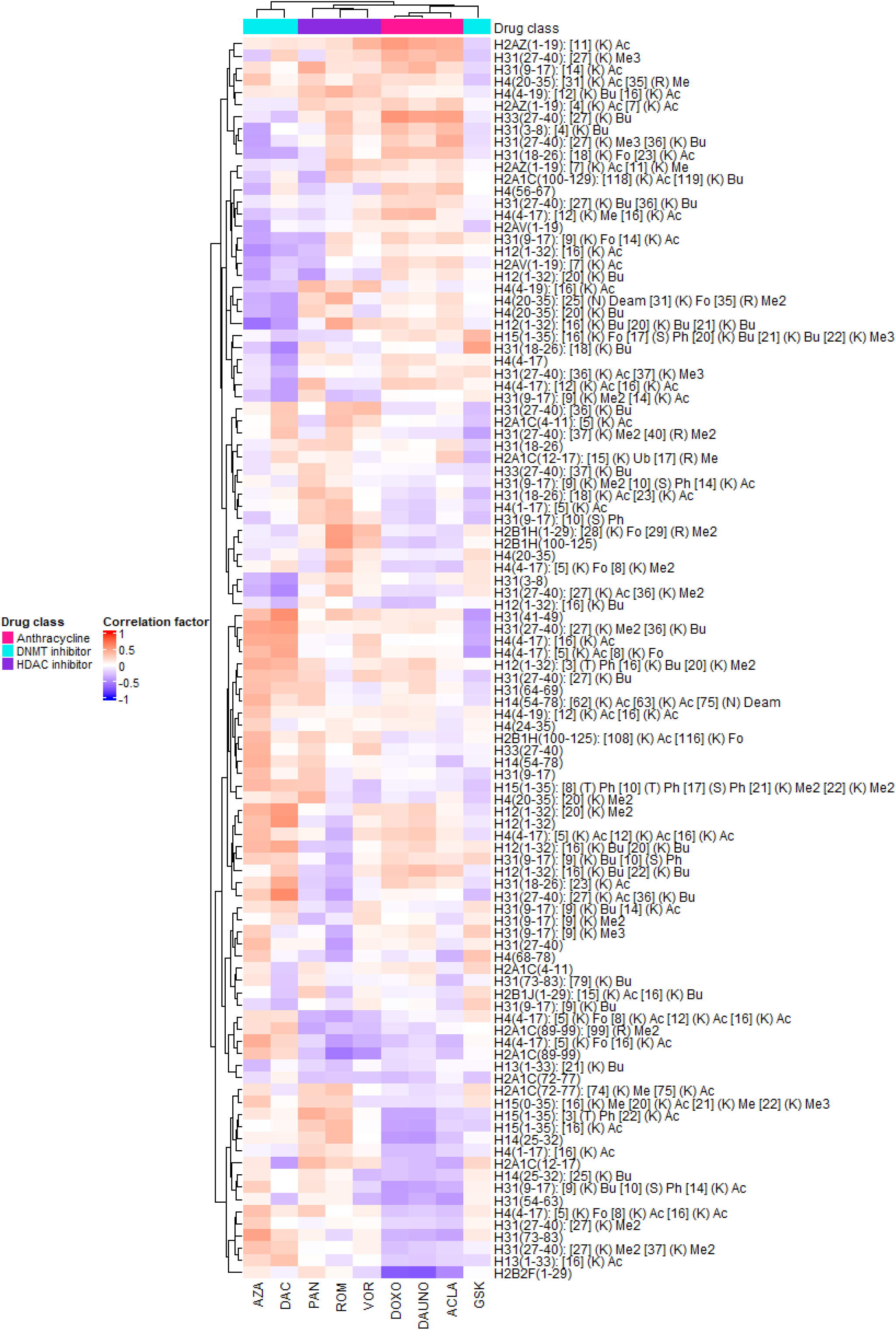
Clustered heatmap showing the Spearman correlation factors between histone peptidoforms (rows) and IC50 values (columns) over the 21 cell lines. Positive correlation coefficients (red) indicate that higher hPTM levels are associated with higher IC50 values, suggesting a link to lower sensitivity. Conversely, negative correlation coefficients (blue) imply that high hPTM levels are associated with lower IC50 values, indicating a potential association to higher sensitivity.

**Supplementary Figure 4.**
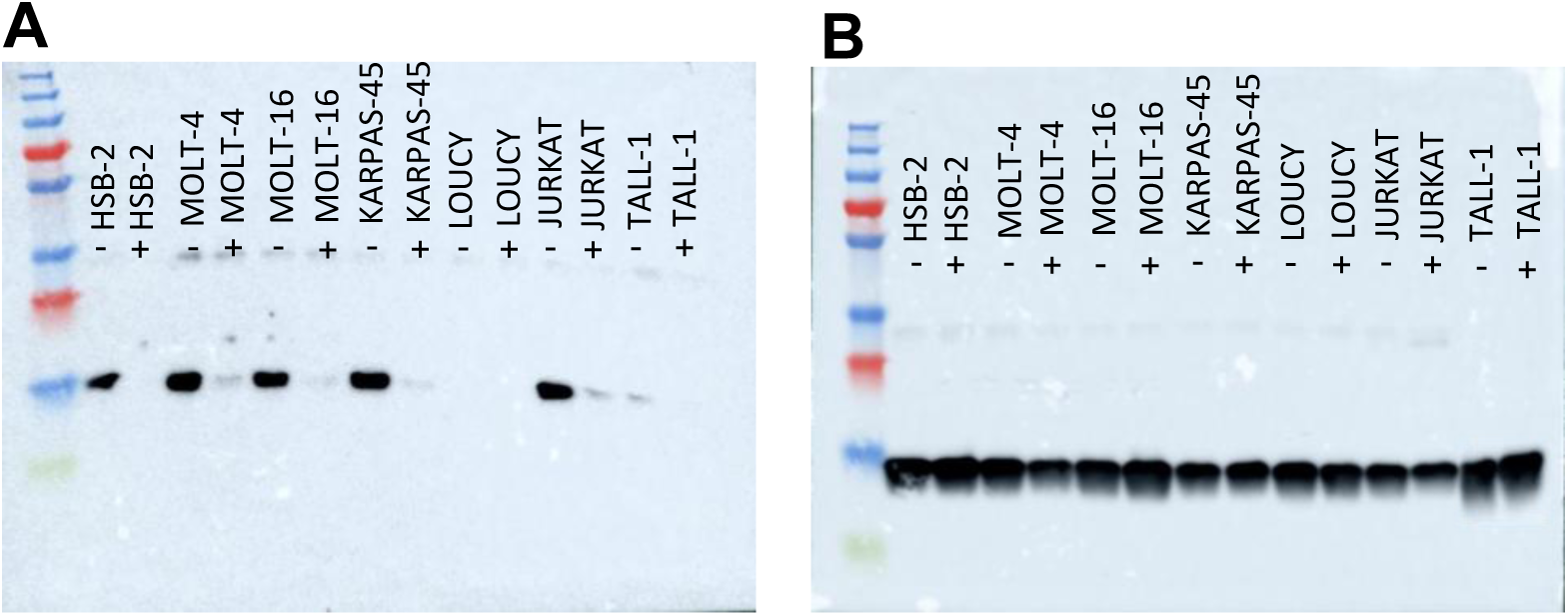
Western blot images showing H3K27me3 **(A)** and H3 **(B)** levels after 24-hour aclarubicin treatment in a subset of seven cell lines with or without 72-hour pre-treatment with the EZH2 inhibitor GSK126.

**Supplementary Figure 5.**
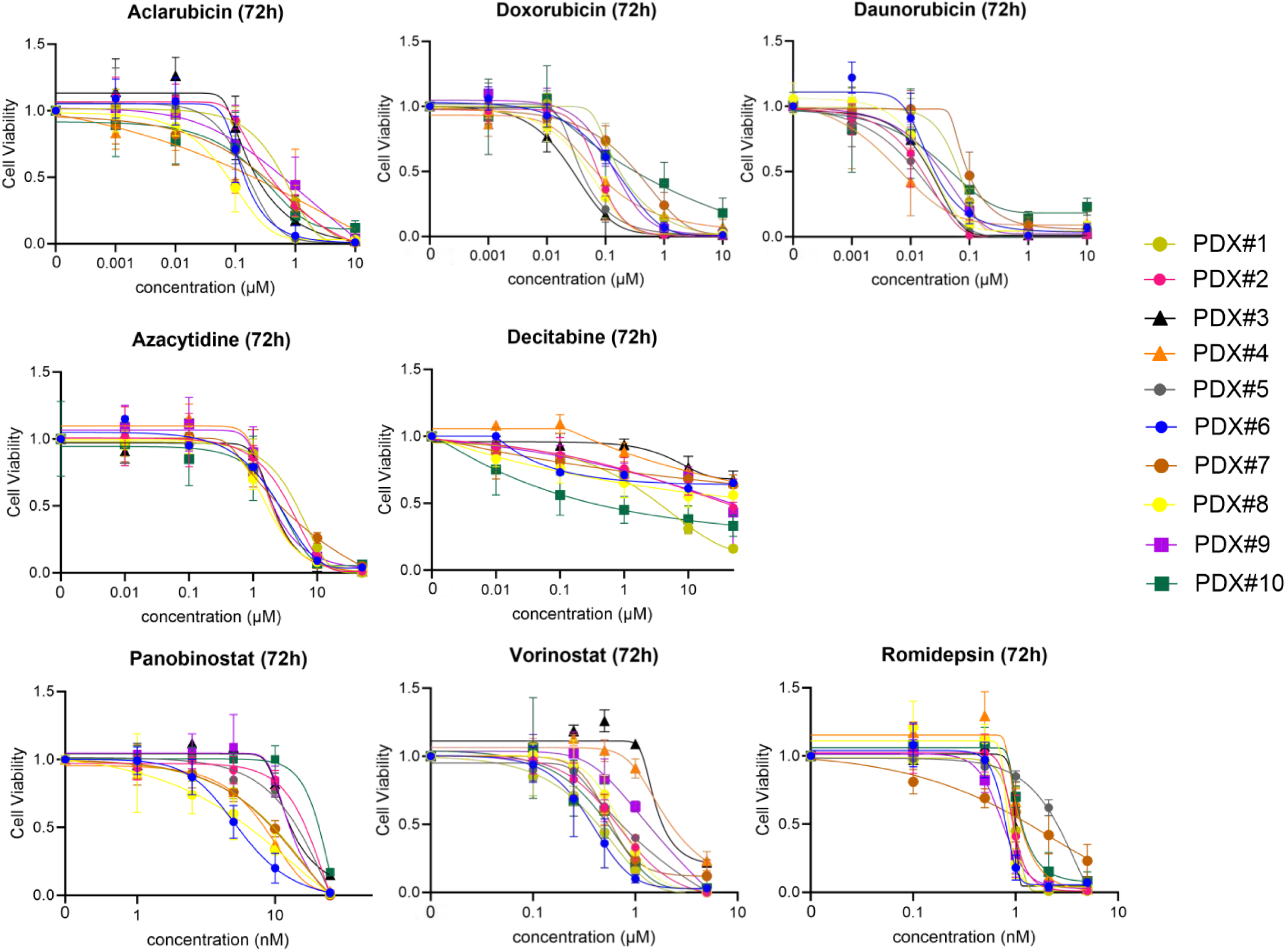
Dose-response curves showing the cell viability of 10 T-ALL PDX models (colors) after in vitro treatment with a dilution series of eight epidrugs. Each data point represents the mean of three independent experiments.

**Supplementary Figure 6.**
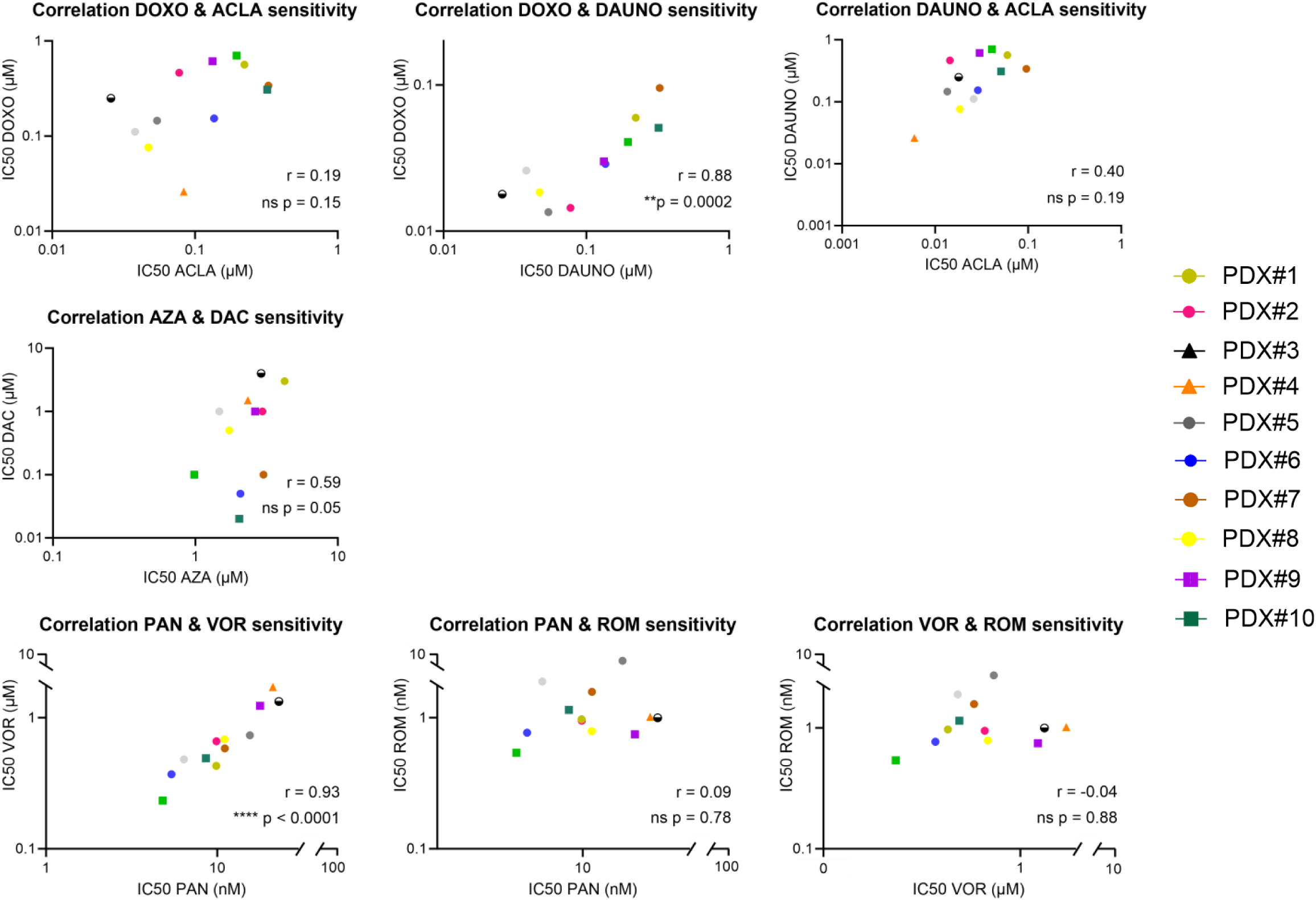
Scatter plots showing the Pearson correlation between the IC50s of 10 T-ALL PDX models within drug classes.

**Supplementary Figure 7.**
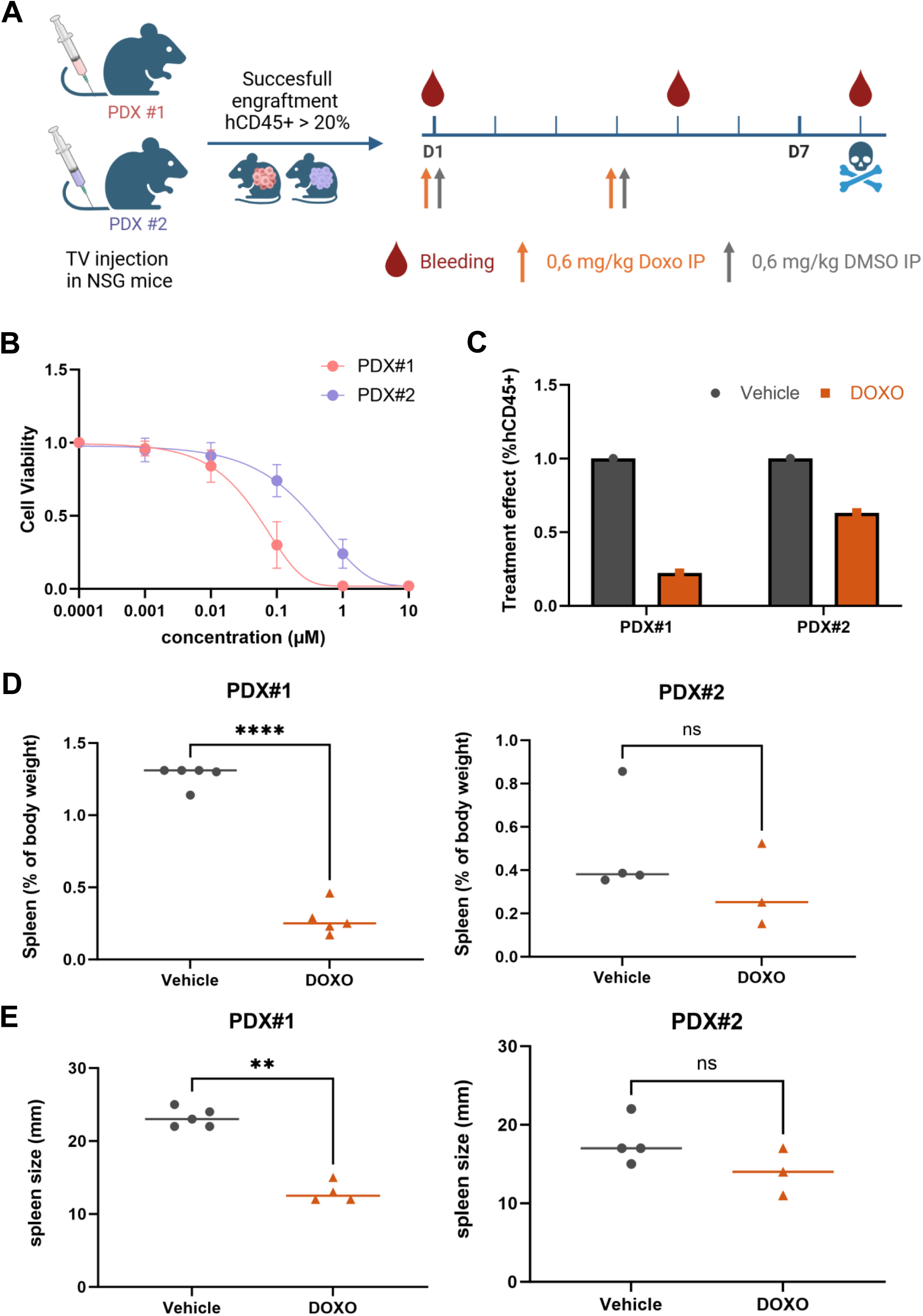
In vivo doxorubicin treatment aligns with ex vivo drug response predictions. **(A)** NSG mice were engrafted with malignant bone marrow material of T-ALL patients. Once engraftment percentage in the blood was higher than 20%, mice were treated with vehicle (DMSO, grey) or doxorubicin (0.6 mg/kg body weight, orange) for two times with an interval of three days. Blood was collected on day one, day five and day eight to assess %hCD45. Mice were sacrificed on day eight for the collection of leukemic blasts. Figure created with BioRender.com. **(B)** Dose-response curve showing the cell viability of both PDX models after ex vivo treatment with a dilution series of doxorubicin. Each data point represents the mean of three independent experiments. PDX#1 is more susceptible to doxorubicin compared to PDX#2. **(C)** The treatment effect of 1 cycle of doxorubicin on hCD45+ cells in the peripheral blood is shown by normalizing tumor growth (average #hCD45+ cells on day 5/average #hCD45+ cells on day 1) in doxorubicin treated mice to tumor growth in vehicle treated mice. **(D)** Spleen weight as a percentage of body weight and **(E) s**pleen size of the mice treated with a vehicle (grey) or doxorubicine (orange) for both PDX models.

**Supplementary Figure 8.**
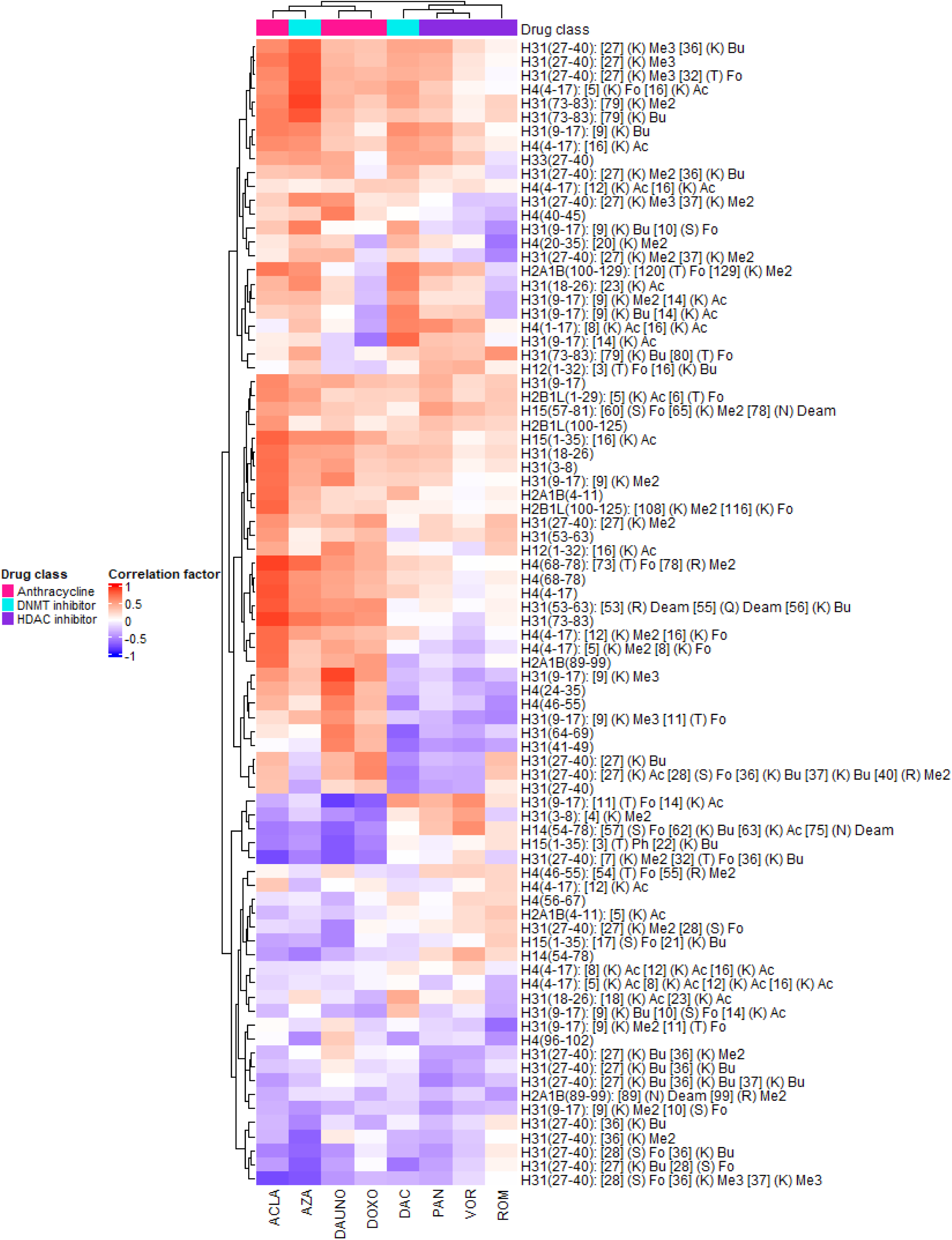
Clustered heatmap showing the Spearman correlation factors between histone peptidoforms (rows) and IC50 values (columns) over the 10 PDX models. Positive correlation coefficients (red) indicate that higher hPTM levels are associated with higher IC50 values, suggesting a link to lower sensitivity. Conversely, negative correlation coefficients (blue) imply that high hPTM levels are associated with lower IC50 values, indicating a potential association to higher sensitivity.

